# Jomon genome sheds light on East Asian population history

**DOI:** 10.1101/579177

**Authors:** Takashi Gakuhari, Shigeki Nakagome, Simon Rasmussen, Morten Allentoft, Takehiro Sato, Thorfinn Korneliussen, Blánaid Ní Chuinneagáin, Hiromi Matsumae, Kae Koganebuchi, Ryan Schmidt, Souichiro Mizushima, Osamu Kondo, Nobuo Shigehara, Minoru Yoneda, Ryosuke Kimura, Hajime Ishida, Yoshiyuki Masuyama, Yasuhiro Yamada, Atsushi Tajima, Hiroki Shibata, Atsushi Toyoda, Toshiyuki Tsurumoto, Tetsuaki Wakebe, Hiromi Shitara, Tsunehiko Hanihara, Eske Willerslev, Martin Sikora, Hiroki Oota

## Abstract

Anatomical modern humans reached East Asia by >40,000 years ago (kya). However, key questions still remain elusive with regard to the route(s) and the number of wave(s) in the dispersal into East Eurasia. Ancient genomes at the edge of East Eurasia may shed light on the detail picture of peopling to East Eurasia. Here, we analyze the whole-genome sequence of a 2.5 kya individual (IK002) characterized with a typical Jomon culture that started in the Japanese archipelago >16 kya. The phylogenetic analyses support multiple waves of migration, with IK002 forming a lineage basal to the rest of the ancient/present-day East Eurasians examined, likely to represent some of the earliest-wave migrants who went north toward East Asia from Southeast Asia. Furthermore, IK002 has the extra genetic affinity with the indigenous Taiwan aborigines, which may support a coastal route of the Jomon-ancestry migration from Southeast Asia to the Japanese archipelago. This study highlight the power of ancient genomics with the isolated population to provide new insights into complex history in East Eurasia.

## Introduction

After the major Out-of-Africa dispersal of *Homo sapiens* around 60 kya, modern humans rapidly expanded across the vast landscapes of Eurasia[1]. Both fossil and ancient genomic evidence suggest that groups ancestrally related to present-day East Asians were present in eastern China by as early as 40 kya[2]. Two major routes for these dispersals have been proposed, either from the northern or southern parts of the Himalaya mountains[1,3–5]. Population genomic studies on present-day humans[6,7] have exclusively supported the southern route origin of East Asian populations. On the other hand, the archaeological record provides strong support for the northern route as the origin of human activity, particularly for the arrival at the Japanese archipelago located at the east end of Eurasian continent. The oldest use of Upper Paleolithic stone tools goes back 38,000 years, and microblades, likely originated from an area around Lake Baikal in Central Siberia, are found in Hokkaido (~25 kya) and mainland Japan (~20kya)[8]. However, few human remains were found from the Upper Paleolithic sites in the Japanese archipelago. Following this period, the Jomon culture started >16 kya, characterized by a hunter-fisher-gathering lifestyle with the earliest use of pottery in the world[9]. Several lines of archaeological evidence support the cultural continuity from the Upper-Paleolithic to the Jomon period, providing a hypothesis that the Jomon people are direct descendants of the Upper-Paleolithic people who likely remained isolated in the archipelago until the end of Last Glacial Maximum[8,10,11]. Therefore, studying the Jomon can provide significant insights into the origin and migration history of East Asians.

A critical challenge for ancient genome analysis in the Japanese archipelago is the inherent nature of warm and humid climate conditions, and the soils indicating strong acidity because of the volcanic islands, which generally result in poor DNA preservation[12,13]. A partial genome of a 3,000-years old Jomon individual from the east-north part of the main-island (Honshu) Japan was reported, but with very limited coverage (~ 0.03-fold) due to the poor preservation[14]. To identify the origin of the Jomon people, we sequenced a 1.85-fold genomic coverage of a 2,500-years old Jomon individual (IK002) excavated from the central part of the Japanese archipelago[15]. Comparing the Jomon whole-genome sequence with ancient Southeast Asians, we previously reported genetic affinity between IK002 and the 8,000-years old Hòabìnhian hunter-gatherer[15]. This direct evidence on the link between the Jomon and Southeast Asians, thus, confirms the southern route origin of East Asians.

Nevertheless, key questions still remain as to (1) whether the Jomon were the direct descendant of the Upper Paleolithic people who were the first migrants into the Japanese archipelago and (2) whether the Jomon, as well as present-day East Asians, retain ancestral relationships with people who took the northern route. Here, we test the deep divergence of the Jomon lineage and the impacts of southern versus northern-route ancestry on the genetic makeup of the Jomon.

## Results

### Ancient DNA prescreening, dating, and sequencing by high-throughput sequencers

Initially, we extracted DNA of twelve individuals from three sites (Fig.S1), which were accompanied by remains associated with the Jomon culture. The endogenous DNA content for ten out of twelve individuals were less than 1.0%, indicating poor preservation of DNA, while those of two individuals, IK002 and HG02, were more than 1.0% (Table S1). From those remaining two individuals, only the ~2,500 years old IK002 (Text S1) exhibited typical ancient DNA damage patterns[16–18](Fig.S2), and was therefore selected for further sequencing. A total of 29 double-stranded sequencing libraries were generated using DNA extracted from the teeth and petrous bone of IK002, yielding endogenous DNA contents ranging from 1.14 to 19.09% (Table S2 &Text S2). The libraries were sequenced to average coverages of 1.85-fold for autosomal genome and 146-fold for mitochondrial genome, with low estimated contamination rate of 0.5% (95% CI: 0.01-2.2%, Fig.S3)[19]. We found IK002 to be assigned to mitochondrial haplogroup N9b1, which is rare among present-day Japaneses people (< 2.0%), but typically found in previous studies of Jomon mtDNA[20,21].

### Testing whether IK002 is the direct descendant of the Upper Paleolithic people in the Japanese archipelago

To make inferences on the genetic relationship of IK002 with geographically diverse human populations, we merged the IK002 genomic data with a diverse panel of previously published ancient genomes[22–28], as well as 300 high-coverage present-day genomes from the Simons Genome Diversity Project (SGDP)[29]. We extracted genotypes for a set of 2,043,687 SNP sites included in the “2240K” capture panel[30]. First, we characterized IK002 in the context of worldwide populations after the out-of-Africa expansion using principal component analysis (PCA)[31,32]. We found that IK002 clusters between present-day Southeast and East Asians and the Upper-Paleolithic human remain (40 kya) from Tiányuán Cave[28,33](Fig.1A). Second, when using a smaller number of SNPs (41,264 SNPs) including the present-day Ainu[34] from Hokkaido (Fig.S1), IK002 clusters with the Hokkaido Ainu (Fig.S4), supporting previous findings that they are direct descendants of the Jomon people[14,34–41]. Thus, the PCA plot showed that IK002 is slightly different from present-day people in East Eurasia and Japan except for the Hokkaido Ainu.

**Fig. 1:**
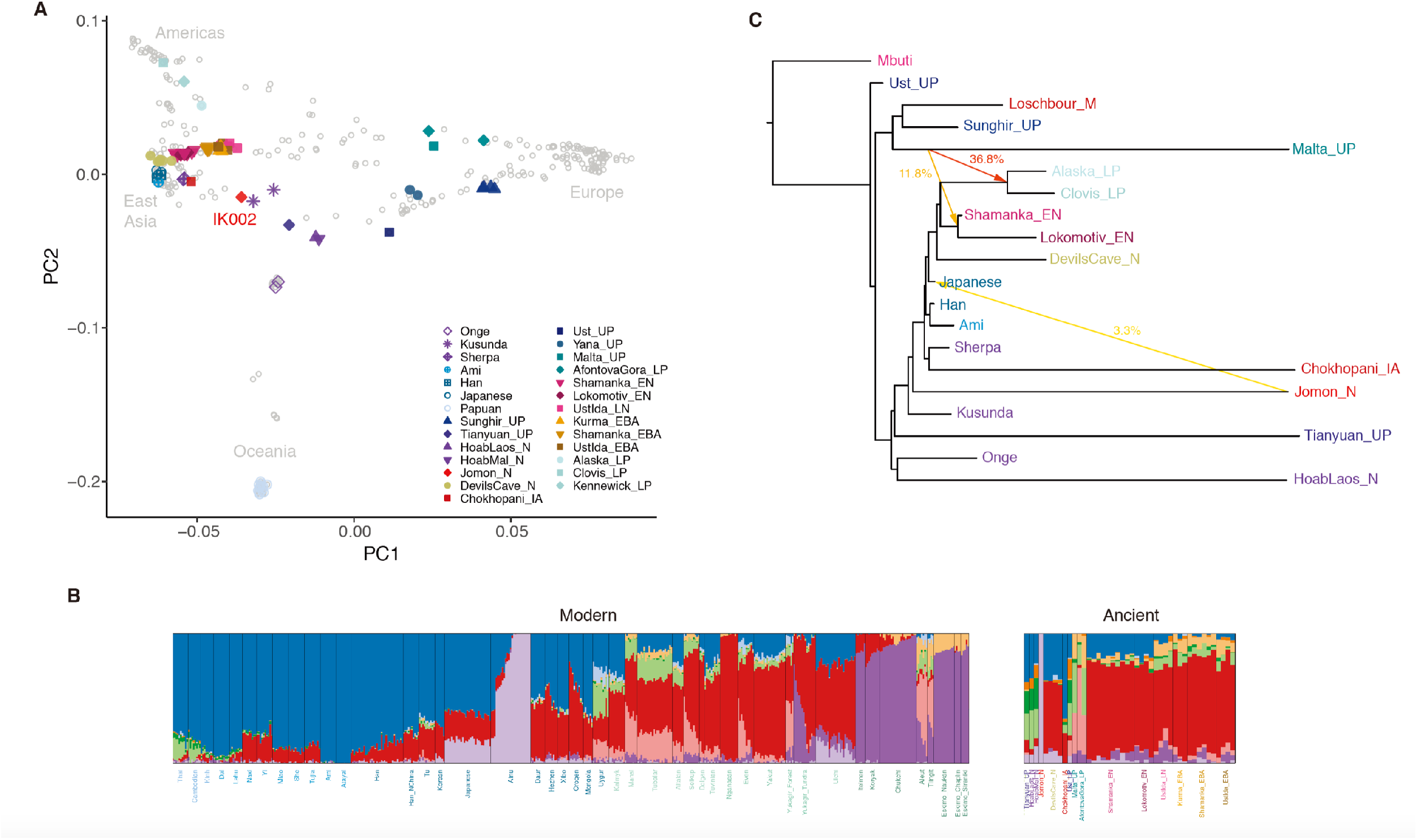
Genetic structure of present-day and ancient Eurasian and Ikawazu Jomon. **A,** Principal component analysis (PCA) of ancient and present-day individuals from worldwide populations after the out-of-Africa expansion. Grey labels represent population codes showing coordinates for individuals. Coloured circles indicate ancient individuals. **B,** ADMIXTURE ancestry components (K=10) for ancient and selected contemporary individuals. The color of light purple represents the component of IK002 which is shared with the present-day Japanese and Ulchi. **C,** Maximum-likelihood phylogenetic tree (*TreeMix*) with bootstrap support of 100% unless indicated otherwise. The tree shows phylogenetic relationship among present-day Southeast/East Asians, Northeast Siberians, Native Americans, and ancient East Eurasians. Mbuti are the present-day Africans; Ust’Ishim is an Upper-Paleolithic individual (45 kya) from Western Siberia (Fu et al., 2014); Mal’ta (MA-1) and Sunghir is Upper-Paleolithic individuals (24 kya and 34 kya) (Raghavan et al., 2014; Sikora et al., 2017), and Loschbour is a Mesolithic individual from Central Siberia (Lazaridis et al., 2014); La368 is a pre-Neolithic Hòabìnhian hunter-gatherer (8.0 kya) in Laos and Önge is the present-day hunter-gatherers in the Andaman island, both of who are from Southeast Asia (McColl et al., 2018); Tiányuán is an Upper-Paleolithic individual (40 kya) in Beijing, Chaina; Kusunda are the present-day minority people in Nepal; Chokhopani is an Iron-age individual (3.0 – 2.4 kya) and Sherpa are the present-day minority people, both of who are in Tibet (Jeong et al., 2016); Han, Ami and main-island Japanese are the present-day East Asians (Mallick et al., 2016); Devils Gate Cave is a Neolithic individual (8.0 kya) in the Primorye region of Northeast Siberia, and Shamanka and Lokomotive are Early-Neolithic individuals (8.0 kya) from Central Siberia, respectively (Damgaard et al. 2018); USR1 and Clovis are late-Paleolithic individuals (11.5 kya and 12.6 kya) in Alaska and Montana, respectively (Moreno-Mayar et al., 2018, Rasumussen et al., 2014). Coloured arrows represent the migration pathways and signals of admixture among all datasets. The migration weight represents the fraction of ancestry derived from the migration edge.

Subsequently, we carried out model-based unsupervised clustering using ADMIXTURE[42] (Fig.S5). Assuming K = 10 ancestral clusters (Fig.1B), an ancestral component unique to IK002 appears, which is the most prevalent in the Hokkaido Ainu (average 79.3%). This component is also shared with present-day mainland Japanese as well as Ulchi (9.8% and 6.0%, respectively) (Fig.1C). Thus, the results of ADMIXTURE showed the strong genetic affinity between IK002 and the Hokkaido Ainu.

We next used ALDER[43] to investigate the timing of admixture in populations with Jomon ancestry. Using IK002 and present-day Han Chinese as source populations, we estimate the admixture in modern Japanese to date to between 42 and 57 generations ago (~1,200 - 1,700 years ago assuming 29 years / generation), consistent with previous estimates[44] (Table S3). For the Ulchi we estimate a more recent timing (8-33 generations ago) consistent with the higher variance in Ikawazu Jomon admixture proportions, albeit with overall weaker statistical support[7]. Finally, using IK002 and present-day Japanese as source populations, we detect very recent (2-4 generations ago) admixture for the Hokkaido Ainu, likely a consequence of still ongoing gene flow between the Hokkaido Ainu and mainland Japanese.

To further explore the deep relationships of Jomon and other Eurasian populations, we used *TreeMix*[45] to reconstruct admixture graphs of IK002 and 18 ancient and present-day Eurasian and Native American groups (Fig.1C & Fig.S6). We found the IK002 lineage placed basal to the divergence between ancient and present-day Tibetans[7,29] and the common ancestor of the remaining ancient/present-day East Eurasians[29,46] and Native Americans[47,48]. These genetic relationships are stable across different numbers of migration incorporated into the analysis. Major gene flow events recovered include the well-documented contribution of MA-1 to the ancestor of Native Americans[47], as well as a contribution of IK002 to present-day mainland Japanese (m=3~6; Fig.S6). IK002 can be modelled as a basal lineage to East Asians, Northeast Asia/East Siberians, and Native Americans, supporting a scenario in which their ancestors arrived through the southern route and migrated from Southeast Asia towards Northeast Asia[6,49]. The divergence of IK002 from the ancestors of continental East Asians therefore likely predates the split time between East Asians and Native Americans, which has been previously estimated at 26 kya[47]. Thus, the *TreeMix* strongly supported that IK002 is the direct descendant of the people who brought the Upper Paleolithic stone tools 38,000 years ago into the Japanese archipelago.

### Testing the impacts of the northern route migration into East Asia

Taking advantage of the earliest divergence of the IK002 (Fig.1C & Fig.S6), we address a question if the Upper-Paleolithic people who took the northern route of the Himalayas mountains to arrive east Eurasia made genetic contribution to populations migrated from Southeast Asia. Under the assumption that MA-1 is a descendant of a northern route wave, we tested gene flow from MA-1 to IK002, as well as to the other ancient and present-day Southeast/East Asians and Northeast Asians/East Siberians by three different forms of *D* statistics: *D*(Mbuti, MA-1; *X*, Ami), *D*(Mbuti, *X*; MA-1, Ami), and *D*(Mbuti, Ami; *X*, MA-1).

The first *D* statistics (shown as red in Fig.2) provides results consistent with previous findings on the prevalence of MA-1 ancestry in the present-day Northeast Asians/East Siberians (*Z* < −3; Table S4)[23], while none of Southeast/East Asians, except for Oroqen, shows a significant deviation from zero. The tree relationships observed in Fig. 1C are confirmed from the other two different forms between Ami and all of the tested populations with some variation that is mostly explained by the MA-1 gene flow (cyan and green in Fig.2, Table S5 & Table S6). Therefore, we concluded that no gene flow from MA-1 to the ancient/present-day Southeast/East Asians including IK002 was detected.

**Fig. 2:**
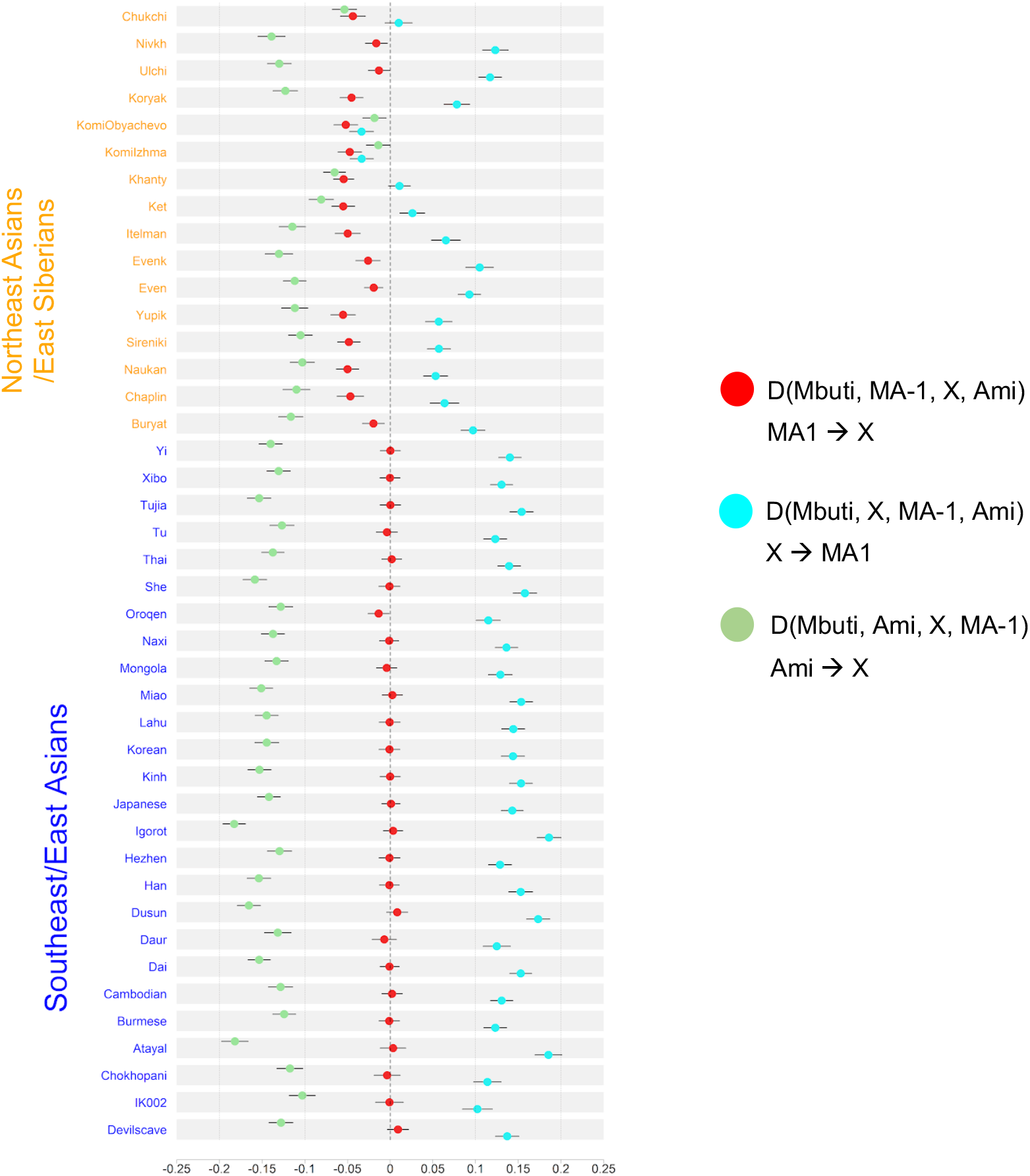
Exploring genetic affinities of IK002 within Northeast Siberians and Southeast/East Asians, respectively. Three different *D* values are plotted with different colors; *D*(Mbuti, MA-1; *X*, Ami) in red, *D*(Mbuti, *X*; MA-1, Ami) in cyan, and *D*(Mbuti, Ami; *X*, MA-1) in green. Error bars show 3 standard deviation, and the vertical dotted and dashed lines indicate *D* = 0 and *D* = −0.2, −0.1, 0.1, and 0.2.

### Remnant Jomon-related ancestry in the coastal regions of East Asia

The basal divergence of IK002 (Fig.1C) suggests a negligible contribution to later ancient and present-day mainland East Asian groups. To further test this prediction we conducted *f*_4_ statistics with the form of (Mbuti, IK002; *X*, Chokhopani). If IK002 were a true outgroup to later East Asian groups, this statistic is expected to be zero for any test population X. However, we find that together with Japanese, present-day Taiwan aborigines (Ami and Atayal), as well as minorities in the Okhotsk-Primorye region (Ulchi and Nivhk) also showed a significant (Z < −3) excess of allele sharing with IK002. Populations in the inland of the eastern part of the Eurasian continent on the other hand were consistent with forming a clade with Chokhopani (Fig.3), suggesting the presence of remnant Ikawazu Jomon-related ancestry in present-day coastal populations in East Asia. The signal is also present in the Neolithic individuals from Devil’s Gate Cave in the Primorye region (Z < −3; Fig.S7a), suggesting that at 8 kya IK002-related ancestry in the region had already been largely but not completely replaced by later migrations. Interestingly, the genetic affinity to IK002 was found, thus, only in the coastal region but not in the inland for both ancient and present-day populations (Fig.3).

**Fig. 3:**
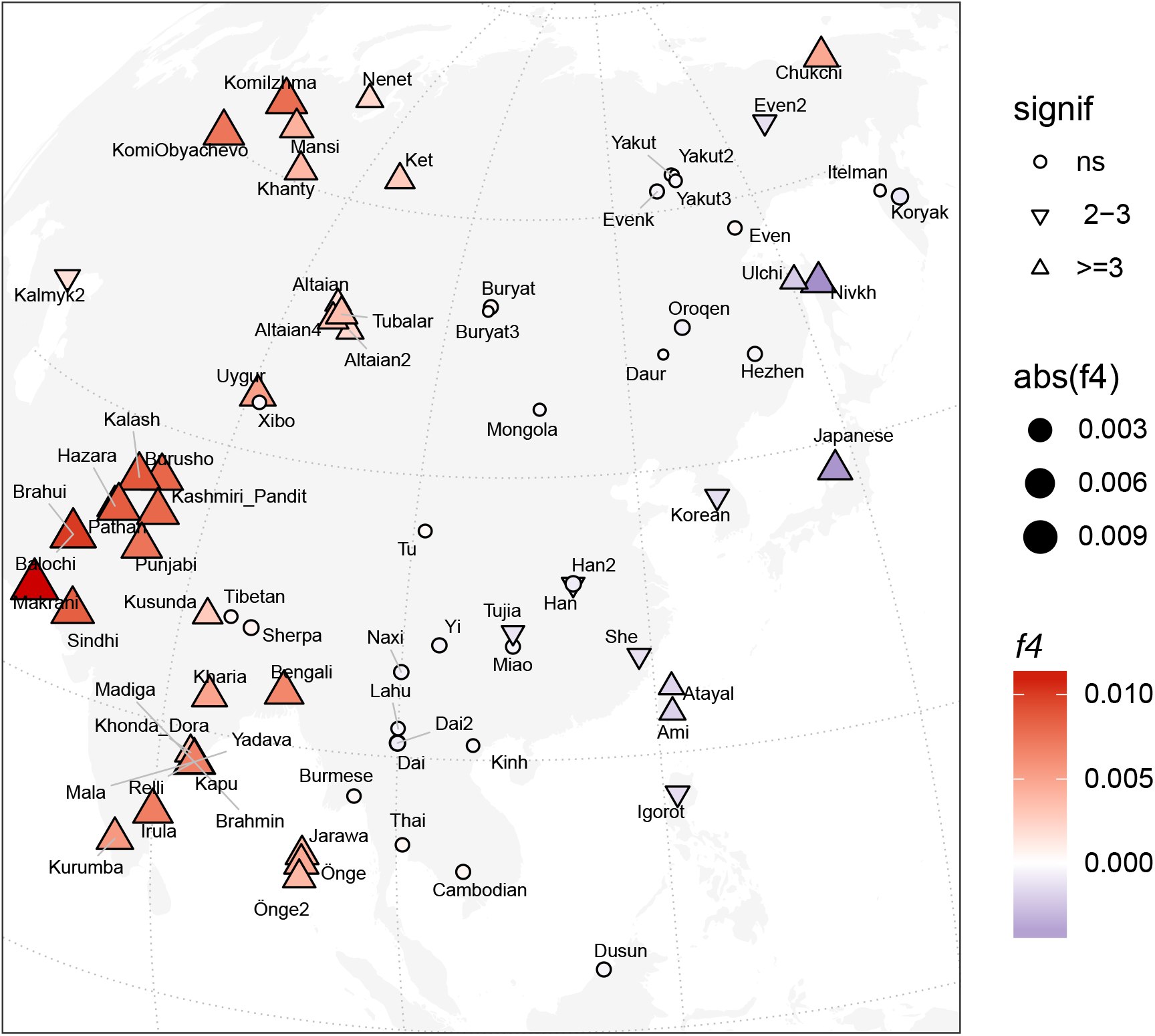
Heatmap of *f*_4_ statistics comparing Eurasian populations to the Ikawazu Individual. Heatmaps of *f*_4_(Mbuti, IK002;*X*, Chokhopani), where *X* are the present-day East Eurasian populations. The colour and size represents the value of *f*_4_ statistics. The shape represents statistical significances of genetic affinities based on Z score. Triangle label means statistical significance with |Z|>3, inverted triangle means weak significance with |Z|=2~3 and circle means non-significance with |Z|<2.

## Discussion

This study takes advantage of whole-genome sequence data from the 2,500-years old Jomon individual, IK002, dissecting the origins of present-day East Asians. IK002 is modelled as a basal lineage to East Asians, Northeast Asians/East Siberians, and Native Americans (basal East Eurasians: bEE)(Fig.4), supporting a scenario in which their ancestors came through south of the Himalayas mountains and migrated from Southeast Asia towards the north[6,49]. We clearly show the early divergence of IK002 from the common ancestor of the other ancient and present-day East Eurasian and Native Americans (Fig.1C). Given that the split between the East Asian lineage and the Northeast Asians/East Siberian and Native American lineage was estimated to be 26 kya[47], the divergence of the lineage leading to IK002 is likely to have occured before this time but after 40 kya when the Tiányuán appeared (Fig.4). Therefore, our results support the archaeological evidence that the Jomon are direct descendants of the Upper-Paleolithic people who started living in the Japanese archipelago 38 kya.

**Fig. 4:**
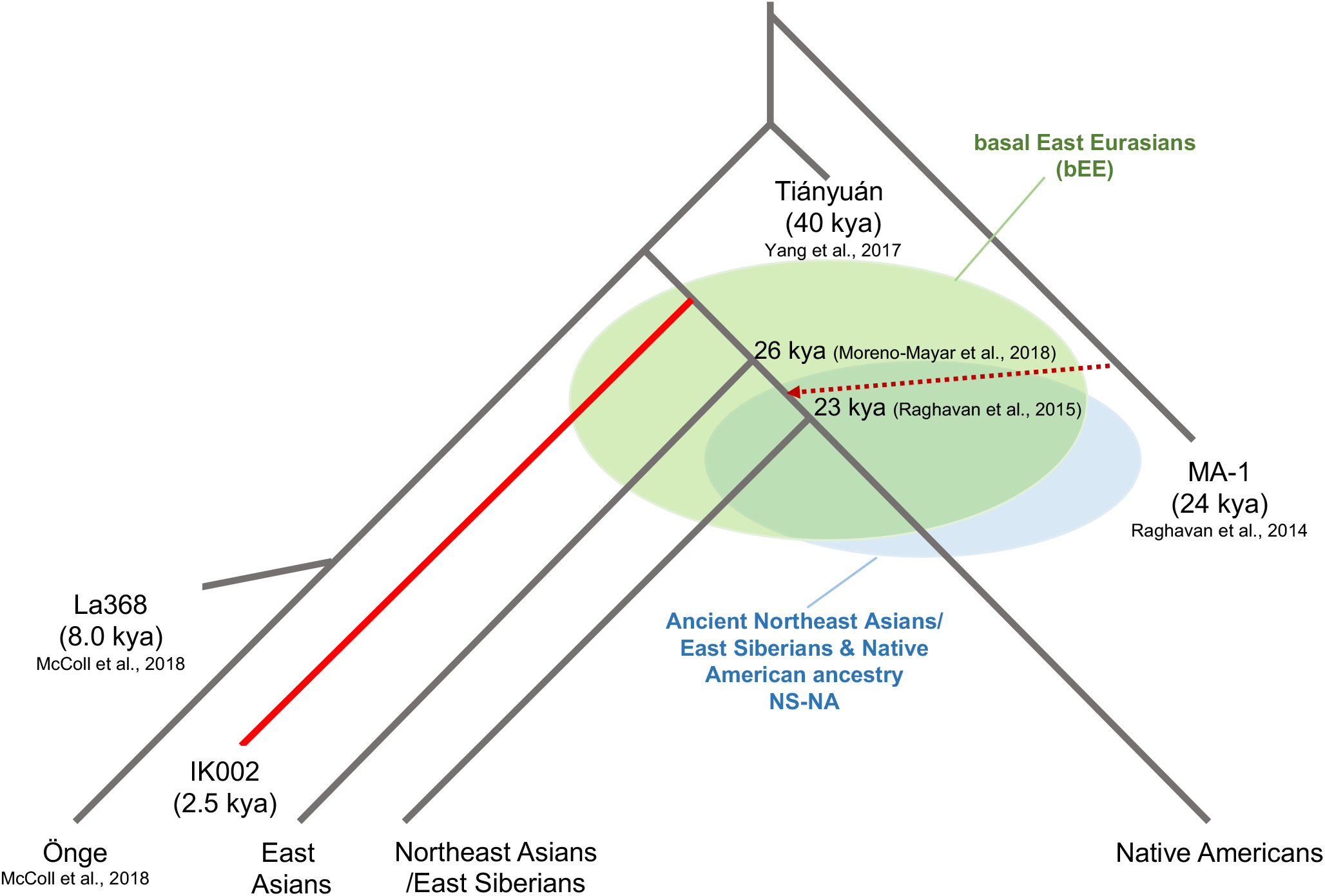
Schematic of peopling history in Southeast and East Asians, Northeast Asian/East Siberians and Native Americans.

Here, we use the MA-1 ancestry as a proxy for ancestral populations who took the northern route of Himalaya mountains to come to East Eurasia. The fine stone tool, *i.e.*, microblade, is a representative technology that was originally developed around Lake Baikal in Central Siberia during the Upper-Paleolithic period[8]. This microblade culture also reached the Hokkaido island ~25 kya and the mainland of the Japanese archipelago ~ 20 kya (Text S5). If this culture was brought by demic diffusion, IK002 may still retain the MA-1 ancestry. However, we find no evidence on the genetic affinity of MA-1 with ancient and present-day Southeast/East Asians including Devil’s Gate Cave (8.0 kya), Chokhopani (3.0 – 2.4 kya), and IK002 (2.5 kya) (Fig.2). Therefore, we conclude that MA-1 gene flow occurred after the divergence between the ancestral populations of Northeast Asians/East Siberians (NS-NA) and East Asians (Fig.4): namely, East Asians originated in Southeast Asia without any detectable genetic influence from the ancestor who took the northern route. There are two hypothetical possibilities to explain the contradiction between genome data and archaeological records. The first possibility is that MA-1 may not be a direct ancestor who invented the microblade culture. The second is that, if the assumption is correct, then the northern-route culture was brought to the Japanese archipelago by the NS-NA population who must have had substantial gene flow from MA-1. The first and second possibilities can be examined by obtaining genome data of the Upper-Paleolithic specimens hopefully accompanying with microblade excavated from around Lake Baikal and from the Primorye region, respectively.

The genome of the Ikawazu Jomon (IK002) strongly supports the classical hypothesis concerning peopling history in the Japanese archipelago. The PCA plot and phylogenetic tree showed that the present-day Japanese fell in the cluster of present-day East Asians (e.g., Han Chinese) but not clustered with IK002 (Fig. 1A & C), while a signal of gene flow was detected from IK002 to present-day Japanese (Fig.1C & S6). The PCA and ADMIXTURE showed the close relationship between IK002 and the Hokkaido Ainu even in the genome-wide structure reflected by linkage blocks (Fig.S4 & 5). These results fit the hypothesis that the Ainu and the Jomon share the common ancestor: the present-day mainland Japanese are the hybrid between the Jomon and migrants from the East Eurasian continent, and the Hokkaido Ainu have less influence of genetic contribution of the migrants[35,40](Text S5).

IK002 gave new insights into the migration route from south to north in East Eurasia. The *f*_4_ statistics suggest that both the ancient and the present-day East Asians are closer to IK002 than Chokhopani (ancient Tibetans, 3.0 – 2.4 kya) in the coastal region but not in the inland region (Fig.3 & Fig.S7). There are two explanations for the genetic affinity to IK002: (1) the earliest-wave of migration from south to north occurred through the coastal region, and/or (2) the migration occurred in both the coastal and inland regions, but the genetic components of the earliest-wave were drowned out by back-migration(s) from north to south occurred in the inland region. In the early migration of anatomically modern humans, the route along the coast has been primarily thought to be important[3,50–53]. The use of water craft could support such explanation for the expansions through the islands and the coastal region[3], which supports the first explanation. There could be, however, potential criticisms: such archaeological evidence of craft boat is hardly found. Ulchi and Nivkh show significantly negative values of *f*_4_ statistics (Z = −4.541 and −10.148, respectively). This could be an influence of the Hokkaido Ainu who are likely to be direct descendants of the Jomon people. The ancestor of the Ainu people could have admixed with the Okhotsk people[54] who were morphologically close related to Ulchi and Nivkh currently inhabit in the the Primorye region[55–57], which are supported by mtDNA[58,59] and genome-wide SNP data (Matsumae et al., unpublished data). The second explanation is that the track of the earliest-wave was erased in the inland but left over in the coastal region. Taiwan aborigines (Ami and Atayal) and Igorot are the Austronesian minorities. Taiwan aborigines are thought to have come from the East Eurasian continent 13.2 +/− 3.8 kya[60], though the origin of Igorot is not well known.

We carried out admixture graph modelling to further characterize the contributions of IK002-related ancestry (Fig. S8). To that end, we first fit a backbone graph including ancient genomes representative of major divergences among East Asian lineages: IK002 (early dispersal); Chokhophani (later dispersal, East Asia) and Shamanka (later dispersal, Siberia). Test populations of interest were then modelled as three-way mixtures of early (IK002) and later (Chokhopani, Shamanka) dispersal lineages, using a grid search of admixture proportions within qpGraph. Consistent with the results from the *f*_4_ statistics, we find that models without contribution from IK002 result in poor model fit scores for Japanese, Devil’s Gate Cave and Ami, as opposed to inland groups such as Han which do not require IK002-related ancestry (Fig. S7). The range of admixture fractions with good model fit is generally quite wide, with best fit models show IK002-related contributions of 8%, 4% and 41% into Japanese, Devil’s Gate Cave and Ami, respectively (Fig. S8). We note that while the substantial contribution into Ami seems at odds with the lower *f*_4_ statistics compared to Japanese, the lineage admixing with Ami shares only a very short branch with IK002, suggesting a contribution from a distinct group with an early divergence from the IK002 lineage. As IK002 also shares ancestry with early Hòabìnhian hunter-gatherers[49], a contribution of those to the ancestors of Ami would also be compatible with this result. However, there is still lack of ancient genome data to understand the peopling history of East Eurasians. It is required to analyze more ancient genome data, if there found appropriate skeletons, in order to fill the gap and to prove the speculation.

## Materials and Methods

### Materials

Twelve individuals from three archaeological sites (Hegi Cave, Hobi and Ikawazu Shell-mounds) were applied to this genomic study. The Hegi Cave site locates on the northern part of Kyushu island, and the Hobi and Ikawazu Shell-mounds, which are close to each other within ~5 km, locate on the center of main island(Honshu) (Fig. S1). Based on conventional chronology of potteries, the individuals from the Hegi Cave site and the Hobi and Ikawazu Shell-mounds were assigned to the early to middle Jomon period (ca. 5,000~8,000 years ago) and the late to final Jomon periods (ca. 3,500~2,500 years ago), respectively (Table S1). Among these Jomon specimens screened in this study, we chose IK002, who were excavated from the Ikawazu Shell-mound site, as the champion sample; we conducted dating IK002 using the bone collagen by AMS, and obtained the calibrated age of 2,720~2,418 calBP (95.4% CI) (Text S1).

### Methods

All the subsequent experiments of ancient DNA were performed in the clean room exclusively built for ancient DNA analyses installed in Department of Anatomy, Kitasato University School of Medicine. The bone and tooth pieces were cut by a sterile disc drill to separate crowns (enamel), roots (cementum and dentine) of teeth for all the samples, and the inner part of petrous bone only for IK002. DNA extraction was carried out with the protocol that is based on the previous studies[22,61] and modified in this study (Text S2). 60 ul DNA extracts obtained from these samples were quantified and qualified by Qubit 3.0 (Thermo Fisher Scientific) and Bioanalyzer (Agilent). Twenty-two extracts were used to construct 34 NGS libraries for Illumina sequencing system using NEBNext Ultra DNA Library Preparation Kit (New England Biolab) in Kitasato University. The NGS sequencing was carried out using MiSeq in Kyushu University and HiSeq in National Institute of Genetics in Japan. For cross-check, we separately prepared another five extracts from IK002, and made the NGS libraries in Copenhagen University and sequenced them using HiSeq in the Danish National High-Throughput DNA Sequencing Centre in Copenhagen (Text S2).

Bioinformatic analyses were carried out in both Kitasato University and Copenhagen University, in order to crosscheck all the data (Text S3 & S4). The contamination rate of IK002 was estimated by ContaMix, and the C-to-T substitution of post-mortem degradation was tested by mapDamage2 (Text S3). To visualize the parametric similarity among Ikawazu Jomon (IK002), the ancient individuals from previous studies, and the present-day human population around the world from the Human Genome Diversity Project (HGDP) and the Simons Genome Diversity Project (SGDP), principal component analyses (PCA) were carried out using EIGENSOFT (Text S4). The genome-wide patterns of population structure and admixture were visualized by ADMIXTURE (Text S4). To reconstruct phylogenetic relationship, TreeMix was used assuming 1 ~ 6 times migration events (Text S4). Pairwise genetic affinities were tested by *f*_4_ and *D* statistics with AdmixTools (Text S4).

## Supporting information

Gakuhari_et_al_2019_SupInfo

## Acknowledgements

We also thank Dr. Yosuke Kaifu for helpful discussion about the idea of two migration routes into East Eurasia. The excavation of the Ikawazu Jomon individual was supported by Grant-in-Aid for Scientific Research (B) (25284157) to YY. The Ikawazu Jomon genome project was organized by HI, and TH & HO who were supported by MEXT KAKENHI Grant Numbers 16H06408 and 17H05132, and by Grant-in-Aid for Challenging Exploratory Research (23657167) and for Scientific Research (B) (17H03738). The Ikawazu Jomon genome sequencing was supported by JSPS KAKENHI Grant Number 16H06279 to ATo, and partly supported in the CHOZEN project in Kanazawa University, and in the Cooperative Research Project Program of the Medical Institute of Bioregulation, Kyushu University. Computations for the Ikawazu Jomon genome were partially performed on the NIG supercomputer at ROIS National Institute of Genetics.

