## Supplementary material for "Jomon genome sheds light on East Asian population history": Gakuhari_et_al_2019_SupInfo

**Supplementary Information**

**Table S1**: Endogenous human DNA content of ancient individuals in prescreening with Miseq.

**Table S2**: Analysis chart of sample ID, position, protocol and laboratory

**Table S3**: Estimation of generation time after admixture between Japanese and Han, Ulchi and Han and Japanese, Ainu and Japanese using ALDER.

**Table S4**: *D* statistics among MA-1, X(the eastern Eurasians) and Ami with Mbuti as the outgroup

**Table S5**: *D* statistics among X(the eastern Eurasians), MA-1 and Ami with Mbuti as the outgroup

**Table S6**: *D* statistics among Ami, X(the eastern Eurasians) and MA-1 with Mbuti as the outgroup

**Figure S1**: Site location of Ikawazu shellmound, Hobi shellmound and Hegi cave in Japanese archipelago.

**Figure S2**: Damage pattern of DNA molecules in a modern human, IK002 and Hegi02 libraries constructed by modified protocols (NEBNext Ultra) in this study. The x-axis means nucleotide position from the end of 5' and 3', and the y-axis means the frequency of substitution for reference human genome (*hg19*). The Blue line represents C-to-T substitution pattern and the red line represents G-to-A substitution. Both frequencies in a modern blood sample is too low. On the other, the frequency of IK002 is clearly higher than that of the modern sample. The frequency of HG02 from the Hegi cave is relatively low than IK002.

**Figure S3**: Evaluating authentication of output data using maximum-likelihood of allele-mismatched patterns with mitochondrial DNA reads using *ContaMix*. **a.**The bootstrap iteration and the frequency of potential contamination risk using mitogenomic reads. The x-axis shows the number of bootstrap iteration and the y-axis shows the shrink factor. The *e* value means the contamination frequency between mitogenomic reads and reference sequences. **b.**The relationship between bootstrap iteration times (x-axis) and probability of estimating precision (y-axis). **c.**The relationship between the probability of estimation (x-axis) and the probability density (y-axis) calculated by the Bayesian estimation.

**Figure S4**: PCA plot including Southeast, East and Northeast Asians, and ancient individuals (IK002, Devil's Gate Cave, Chokhopani and Tianyuan). The small legends with different shapes and colors represent individuals of each populations, and the three-letter abbreviations of population names represent the mean position within each populations.

**Figure S5**: ADMIXTURE ancestry components for ancient and selected contemporary individuals and ancient individuals. The x-axis represents individual data from different populations and the y-axis shows the frequency calculated by haplotype frequencies based on genome-wide SNPs. The color represents the unique component based on assumption of the number of cluster. The Jomon component (purple color) was occurred in K=9~10.

**Figure S6**: TreeMix analysis including Southeast, East and Northeast Asians, Native Americans, and ancient Eurasians, with different migration factors (*m* = 1, 2, 3, 4, 5, 6). The scale bar shows the average standard error (SE) of the entries in the covariance matrix. The scale bar shows the average standard error (SE) of the entries in the covariance matrix. Coloured arrows represent the migration pathways and signals of admixture among all datasets. The migration weight represents the fraction of ancestry derived from the migration edge.

**Figure S7**: Heatmap of $f_4$ statistics comparing eastern Eurasian populations to IK002. Heatmaps of $f_4$(Mbuti, IK002; *X*, Chokhopani), where X are (**a**) modern and (**b**) ancient East Eurasian populations.

**Figure S8**: the estimated admixture models of each populations (Japanese, Devils Cave and Ami) using qpGraph based on three-way mixtures of early (IK002) and later (Chokhopani, Shamanka) dispersal lineages.

**Text S1: Archaeological information and radiocarbon dating of IK001 and IK002**

The Ikawazu Shell-mound site locates in the Tahara city [34° 38′ 43″ north latitude; 137° 8′ 52″east longitude], Aichi Prefecture, where is in the central part of main island of the Japanese archipelago. The Ikawazu Shell-mound was initially excavated in 1918[1]: over 100 individuals were excavated from the site, accompanied with the Jomon potteries assigned to the late - final Jomon period (ca. 3,500-2,500 years ago) based on the pottery chronology. More recently, one of us (Y.M.) and colleagues excavated the new section within the Ikawazu Shell-mound site in 2010 and found six buried individuals of complete skeletal remains. IK001 and IK002 were recovered from Pit No.4, and showed a better state of preservation than those of the other remains. IK001 and IK002 had morphologically typical Jomon characteristics. On the side of the IK002 head, a Jomon pottery, so called a Gokan-no-mori type Jomon ware-corded which is typical in the late Jomon period, was offered. IK001 was excavated together with IK002: the former was an infant, and the latter was a late-middle-age woman. Preliminary PCR-direct sequencing of mitochondrial DNA (mtDNA) showed IK001 and IK002 had different mt D-loop sequences, suggesting they were not a mother-child relationship.

We extracted the collagen from IK001 and IK002, and obtained the purified gelatins for radiocarbon dating that were carried out using a compact AMS at The University Museum in The University of Tokyo. The conventional radiocarbon ages were estimated to 2638±16 BP and 2681±16 BP, respectively. Given that the Ikawazu people built the large shell-mound, it is likely that IK001 and IK002 had applied to marine resources. To correct the marine reservoir effect depending on intake ratio of marine fish and shell, we measured stable isotope ratios of carbon and nitrogen from extracted gelatins of IK001 and IK002. Calculating contribution of amino acids from marine resources with the two end-points of terrestrial herbivore and marine fish, the 50 % marine were estimated. These data were calibrated by OxCal 4.2 (calibration program based on the calibration curve of IntCal 13), and the calibrated ages of IK001 and IK002 showed 2,699~2,367 cal BP (95% CI) and 2,720~2,418 cal BP (95.4% CI), respectively. Because these ages were assigned to the Gokan-no-mori period which has no evidence of rice cultivation, we confirmed that IK001 and IK002 were individuals from the Jomon period accompanied with typical Jomon culture. We sampled teeth of IK001 (M1) and IK002 (M3), and fragments of the petrous bone of IK002.

**Text S2: DNA extraction and library construction**

The tooth samples were cut by a sterile and UV-irradiated disc drill to separate crown (enamel) and root (cementum and dentine). DNA extraction of the root was carried out by the Gamba method[2,3]with our modification. The teeth were washed by 3% sodium hypochlorite solution (Sigma-Aldrich) for 15 minutes, in order to decrease the degree of modern DNA contamination. After washing the teeth with ultrapure water (Thermo Fisher Scientific) and 99% ethanol (Sigma-Aldrich), the teeth were dried on the clean bench in the clean room for 16 hrs. The washed samples were pulverized by freezer mill (ShakeMaster Auto ver 2.0, BioMedical Science Inc.), and fine powder was obtained.

To release DNA molecules from the sample powder, 200 mg tooth powder was incubated for 24 hrs at 55℃ followed by 24 hrs at 37℃in 2 ml DNA LoBind tube (Eppendorf) with 1ml lysis buffer in final concentrations of 20mM Tris HCl (pH 7.5), 0.7% N-lauroylsarcosine, 47.5mM EDTA (pH 8), 0.65U/ml Proteinase K, shaking at 900 rpm in a Thermomixer (Eppendorf). The samples were then centrifuged at 13,000 g for 10 min and the supernatants were discarded. Fresh lysis buffer (1 ml) was added to the pellet, vortexed, and the incubation and centrifugation steps were repeated. The second supernatants were then transferred to ultrafiltration tubes (Amicon® Ultra-4 Centrifugal Filter Unit 30K, Merck), diluted with 3ml TE (pH 8.0) and centrifuged at 2,000 g until final concentrations of ~100 mlwere obtained. These volumes were then transferred to silica column (MiniElute PCR Purification Kit, QIAGEN) and purified according to manufacture's instructions, except for adding TWEEN 20 (at 0.05% final concentration) to 60 ml EB buffer pre-heated to 60℃ at the final step.

The petrous bone was cut by a sterile and UV-irradiated disc drill, and three pieces where Pinhasi et al. (2015) named as "C-part"[4] were obtained (C1, C2, C3); the pieces were washed by ultrapure water (Thermo Fisher Scientific) and 99% ethanol (Sigma-Aldrich). After the dried pieces were drilled and homogenized, ~500 mg bone powder was obtained from the three pieces. The first powder of 150 mg was used to extract DNA molecules following the modified protocol mentioned above. The powder of C2 was rinsed by ultrapure water [Rinsed supernatant], then treated with pre-digestion buffer containing 20mM Tris HCl (pH 7.5), 0.7% N-lauroylsarcosine, 0.4M EDTA (pH 8), 0.65U/ml recombinant Proteinase K for 30 min at shaking at 900 rpm in a Thermomixer (Eppendorf). The mixture was then centrifuged at 13,000 g for 10 min and the supernatant was transferred to a 2 ml tube DNA LoBind [Pre-digestion]. Fresh lysis buffer (1 ml) containing 20mM Tris HCl (pH 7.5), 0.7% N-lauroylsarcosine, 47.5mM EDTA (pH 8), 0.65U/ml recombinant Proteinase K was added to the pellet. After vortexed and incubated for 24 hrs at 55℃ followed by shaking at 900 rpm for 24 hrs at 37℃, the first extract was obtained [Extract 1]. This step was then repeated, and the second extract [Extract 2] was obtained. The residual pellet was pulverized by wet-grinding with shaking sterile beads in grinding cylinder. Fresh lysis buffer containing 20mM Tris HCl (pH 7.5), 0.7% N-lauroylsarcosine, 0.4M EDTA (pH 8), 0.65U/ml recombinant Proteinase K was added into the pulverized pellet, and the pellet was incubated for 24 hrs at 55℃ followed by shaking at 900 rpm for 24 hrs at 37℃ in 2 ml tube, the third extract was obtained [Extract 3]. The five elutes (rinsed and pre-digestion supernatants and three extracts)were filtrated following the protocol mentioned in the paragraph of DNA extraction from tooth. Finally, we obtained four DNA extracts from each petrous bone piece (total 15 extracts). We used [Extract 2] of C1 and C3 to construct NGS libraries.

Two different protocols of construction NGS libraries from five elutes of C2 ([Rinsed supernatant], [Pre-digestion], [Extract 1], [Extract 2], [Extract 3]) were used separately in two laboratories (Kitasato University School of Medicine in Japan, and Copenhagen University Geogenetics Laboratory in Denmark) for inter-laboratory crosschecking. In the Kitasato University, the bead-based size selection protocol with NEBNext Ultra DNA library preparation kit (New England Biolabs: NEB) was used. To remove large DNA fragments that could be contaminants from modern organisms, we modified the NEB original protocol: we adjusted the mixing ratio of the Agencourt AMPure XP solution (Beckman Coulter), the Solid Phase Reversible Immobilization (SPRI) magnetic bead solution, for DNA fragment solution. In the Copenhagen University, the protocol shown in Allentoft et al. (2015)[5] was used to make NGS libraries. Eventually, we constructed 6 libraries from tooth and 18 libraries from the petrous bone in the Kitasato University, and 5 libraries from the petrous bone in the Copenhagen University; totally we provided 29 libraries from IK002.

**Text S3:Data output, processing, and authentication**

The 29 libraries were sequenced on a flowcell using the Ilumina HiSeq 2500 and the HiSeq reagent kit of normal and rapid mode for 100 cycles in paired end in the National Institute of Genetics in Japan and the Danish National High-Throughput DNA Sequencing Centre in Denmark. After running HiSeq, the sequence reads were called by Illumina's Real Time Analysis (RTA) or CASAVA 1.8.2 (Illumina) base-calling software. The HiSeq output-data were processed using customizable NGS pipeline in the Geogenetics Laboratory and the Kitasato University. AdapterRemoval v. 2[6] was used to trim adapters terminal N's (--trimns), low quality bases (-trim qualities, --minquality 2) and short reads (--minlength 30), and filtered reads were checked with FastQC v. 0.11.7[7]. The filtered reads were mapped against *hg19*, human reference genome, by BWA ver. 0.5.9. Mapped reads with mapping quality below Phred score 30 and duplicates were removed using SAMtools[8] and the MarkDuplicates tool of Picard Tools (http://broadinstitute.github.io/picard/ ). Read depth and coverage were determined using pysam and BEDtools.

Misincorporation patterns were assessed using mapDamage2[9]. The degree of modern DNA contamination was estimated by *ContamMix[10]*focused on mitogenome sequences. The resulting sequence assembly and haplotype for mitochondrial genome was visualized using MitoSuite v.1.0.9[11]. After sorting, indexing and -mpilup the mapped reads using Samtools -sort and -index and vcftools, SNP calling and extracting on nuclear genome against 2240K panel were carried out by GATK-2.2-3[12]and PLINK v1.9[13].

**Text S4:Population genetic analyses**

Principal component analysis

As a first assessment of the genetic affinities of the study individuals we carried out principal component analysis (PCA), as previously described[14–16]. In particular, we projected the low coverage ancient individuals onto the PCs inferred from different sets of modern and high coverage ancient individuals, using the 'lsqproject' option in smartpca from the EIGENSOFT package[14]. To reduce the influence of C > T and G > A substitutions at the both end of DNA molecules, we extracted SNPs from transversion sites in PLINK datasets.

ADMIXTURE

To explore sheared genetic component between IK002 and the other populations, we ran ADMIXTURE v1.3.0[17]from K=1 to K10 on the SGDP panel, and the Ainu people in present-day populations and the Chokhopani 1 and Tiányuán individuals in ancient populations, selecting the best of 500 replicate runs for each value of K with LD-pruned datasets (block size = 5 Mb). Genotypes where the ancient individuals showed the damage allele at C > T and G > A SNPs were excluded for each low coverage ancient individual.

*TreeMix*

The datasets for *TreeMix*[18], the program based on haplotypes of each block punctuated on human genome with LD-pruning definition, were chosen to represent different ancestries of East Eurasians and Native Americans; IK002, East Asians (Han, Ami, Japanese and Devils Cave), Northeast Siberians (Lokomotiv and Shamanka, the ancient Siberians), Native Americans (Clovis and USR1, the ancestry of Native American), Himalayan (Sherpa, Kusunda and Chokhopani, the ancient highlander) and Southeast Asians (Önge and La368, the Hoabinhian). Furthermore, Tiányuán, Mal'ta (MA-1) and Ust'Ism were included as a landmark of divergence events happened in the Upper Paleolithic period. We used Mbuti as an outgroup and ran different conditions with regard to migration events ($m$= 0 ~ 6) allowed to fit admixture graphs to the data. We only considered the SNP sites that are non-missing in all individuals included in this analysis and chose the tree under each condition that showed the highest likelihood among 1,000 trees with different random seeds.

*f*-statistics and *D*-statistics

We used the *D*statistic framework and $f_4$statistical analyses to investigate patterns of admixture and shared ancestry in our data set. All *D*statistics were calculated from allele frequencies using the estimators described previously[5,15], with standard errors obtained from a block jackknife (the jackknife parameter = 0.050, the number of blocks = 714). Calculating of *D*statistics was carried out by *qpDstat* in the AdmixTools 4.1. The values of *D*statistics were visualized and mapped by *R*software. $f_4$statistics was calculated by *qpDstat* with the $f_4$ mode.

**Text S5:Geological history, archeological/anthropological records, and history of studies on peopling history in the Japanese archipelago**

The trace of *Homo sapiens* activity in Japan appears around 38 kya when the Japanese archipelago was connected with the Eurasian continent (Maritime Province of Siberia) by land through the current Sakhalin and Hokkaido islands (so called, the Paleo-SHK peninsula). Because the Blakiston line (the Strait of Tsugaru) between the Paleo-SHK peninsula island and the Paleo-Honshu island (current main-island) was much shorter than present state (Fig. S1), it was likely to cross easily the narrow strait for the migrants from the East Eurasia continent (Hokkaido Route). At the time, the Kyushu island was not connected with the Korean peninsula. In the coldest period of Last Glacial Stage (LGS: ~20 kya), the sea level went down, and the Straits of Korean and Tsushima were considerably narrowed (Fig. S1). Then, another route was opened for the migrants into the northern part of Kyushu island through the Korean peninsula (Tsushima Route). Meanwhile, the Ryukyu islands were not connected between the Taiwan island and the Kyushu island at the time (Fig. S1), though the Taiwan island was connected with the southern China. If migrants used boats, they could have reached the southern part of Kyushu island (Okinawa Route). These three were possible routes in the Upper Paleolithic period from East Eurasia into the region of the current Japanese archipelago [19].

Few skeletal remains reliable have not been excavated from the Paleolithic sites in the Japanese archipelago, but from the Ryukyu islands[20,21]. Sato et al. (2015) suggested, however, that no genetic contribution from such Upper Paleolithic people directly to the present-day inhabitants was detected based on modern genomic data from the Ryukyu islands[22]. After the end of LGS, the Japanese archipelago has been separated completely from the Eurasian continent around 15 - 10 kya;

the Upper Paleolithic people must have been isolated genetically from the other East Eurasian populations until the agricultural migrants come from the continent around 3,000 – 2,000 years ago[23], which is so called the Yayoi period) .

In contrast to the Paleolithic period, a considerable number of skeletal remains have been excavated from the main island (Honshu) in the Jomon period, starts around 16.0 kya in the appearance of the Jomon earthenware and ends around 3 - 2.5 kya when the "end" is defined by staring rice cultivation. The Jomon period corresponds to the Neolithic period in Europe, but the Jomon people did not have large-scale agriculture, but lived in hunting-fishing-gathering. In the Ikawazu Shell-mound site, the people did not start rice cultivation yet, and had typical Jomon culture. Because the cultural continuity has been detected from the Upper Paleolithic to the Jomon, it is usually thought that the Jomon people are direct descendants of the Upper Paleolithic people. In the eastern part of Eurasian continent, frequent migrations and admixture occurred during the Neolithic period. But, in the Japanese archipelago, the Jomon people were isolated because of the geographical condition, and must have had very less influence of such genetic disturbance. Therefore, the Jomon genome has been thought to be not changed a lot from the Upper Paleolithic genome.
