## Supplementary_Tables&Figures for "Jomon genome sheds light on East Asian population history"

### Table S1

| **Sample Name** | **Site name** | **Library Conc. (pM)** | **Total reads** | **Mapped reads (*hg19*)** | **Mapped %** | **C>T %** |
|---|---|---|---|---|---|---|
| IK001 | Ikawazu | 12,229 | 410053 | 594 | **0.14** | NA |
| IK002 | Ikawazu | 5,990 | 333597 | 8324 | **2.50** | >10 |
| HB234 | Hobi | 22,815 | 167190 | 19 | **0.01** | NA |
| HB1123 | Hobi | 26,802 | 371468 | 247 | **0.07** | NA |
| HB40 | Hobi | 22,174 | 387621 | 75 | **0.02** | NA |
| HB828 | Hobi | 27,450 | 429742 | 558 | **0.13** | NA |
| HB952 | Hobi | 29,380 | 236487 | 117 | **0.05** | NA |
| HB1284 | Hobi | 22,069 | 288319 | 662 | **0.23** | NA |
| HG02 | Hegi | 17,796 | 408947 | 6536 | **1.60** | <0.5 |
| HG16 | Hegi | 28,746 | 520340 | 962 | **0.18** | NA |
| HG29 | Hegi | 21,433 | 355466 | 181 | **0.05** | NA |
| HG55 | Hegi | 8,935 | 468529 | 2471 | **0.53** | **NA** |

### Table S2

| Sample | Lib | TotalLibReads | TrimLibReads | MappedLibReads | Duplicates | FinalLibReads | Duplicates% | Endogenous% | Efficiency% | Position | Library construction | Institute name of NGS data output |
|---|---|---|---|---|---|---|---|---|---|---|---|---|
| Jomon | Lib01 | 397,457,388 | 205,417,156 | 3,575,342 | 2,897,329 | 678,013 | 81.04 | 1.74 | 0.17 | | | |
| Jomon | Lib02 | 483,234,318 | 249,273,335 | 4,405,539 | 3,713,232 | 692,307 | 84.29 | 1.77 | 0.14 | | | |
| Jomon | Lib03 | 529,297,850 | 269,845,196 | 3,460,131 | 2,115,060 | 1,345,071 | 61.13 | 1.28 | 0.25 | Tooth dentine | NEBNext Ultra DNA library kit | NIG, Mishima, 4~25 Mar., 2016 |
| Jomon | Lib04 | 240,478,394 | 124,698,389 | 2,126,996 | 1,831,771 | 295,225 | 86.12 | 1.71 | 0.12 | | | |
| Jomon | Lib05 | 236,015,856 | 122,502,006 | 2,104,751 | 1,810,212 | 294,539 | 86.01 | 1.72 | 0.12 | | | |
| Jomon | Lib6 | 433,647,102 | 221,339,926 | 2,817,039 | 1,580,164 | 1,236,875 | 56.09 | 1.27 | 0.29 | | | |
| Jomon | I04 | 237,971,558 | 122,881,163 | 7,385,634 | 5,524,508 | 1,861,126 | 74.80 | 6.01 | 0.78 | | | |
| Jomon | I05 | 227,075,442 | 117,476,528 | 7,330,363 | 5,426,615 | 1,903,748 | 74.03 | 6.24 | 0.84 | | | |
| Jomon | I06 | 231,544,700 | 121,574,515 | 6,851,193 | 4,664,805 | 2,186,388 | 68.09 | 5.64 | 0.94 | | | |
| Jomon | I07 | 215,787,182 | 113,748,656 | 6,303,999 | 4,553,925 | 1,750,074 | 72.24 | 5.54 | 0.81 | | | |
| Jomon | I09 | 237,651,346 | 115,421,561 | 3,336,984 | 212,551 | 3,124,433 | 6.37 | 2.89 | 1.31 | | | |
| Jomon | I10 | 237,907,508 | 119,164,747 | 19,111,803 | 2,447,870 | 16,663,933 | 12.81 | 16.04 | 7.00 | | | |
| Jomon | I13 | 234,657,460 | 120,814,912 | 7,476,311 | 5,404,946 | 2,071,365 | 72.29 | 6.19 | 0.88 | | | |
| Jomon | I14 | 235,799,456 | 117,649,335 | 8,331,258 | 6,443,033 | 1,888,225 | 77.34 | 7.08 | 0.80 | | | |
| Jomon | I15 | 235,981,678 | 116,175,678 | 8,661,251 | 6,764,525 | 1,896,726 | 78.10 | 7.46 | 0.80 | Petrous bone | NEBNext Ultra DNA library kit | NIG, Mishima, 20 Feb., 2017 |
| Jomon | I16 | 233,981,444 | 117,490,393 | 2,162,154 | 1,918,880 | 243,274 | 88.75 | 1.84 | 0.10 | | | |
| Jomon | I18 | 235,630,834 | 121,236,313 | 1,973,621 | 1,741,222 | 232,399 | 88.22 | 1.63 | 0.10 | | | |
| Jomon | I19 | 232,848,604 | 119,348,059 | 1,981,256 | 1,744,192 | 237,064 | 88.03 | 1.66 | 0.10 | | | |
| Jomon | I20 | 230,906,338 | 118,884,804 | 1,942,120 | 1,681,914 | 260,206 | 86.60 | 1.63 | 0.11 | | | |
| Jomon | I21 | 234,290,444 | 119,349,954 | 2,017,349 | 1,757,562 | 259,787 | 87.12 | 1.69 | 0.11 | | | |
| Jomon | I22 | 230,406,198 | 119,300,898 | 1,962,062 | 1,694,754 | 267,308 | 86.38 | 1.64 | 0.12 | | | |
| Jomon | I23 | 232,333,464 | 118,067,281 | 2,026,813 | 1,785,558 | 241,255 | 88.10 | 1.72 | 0.10 | | | |
| Jomon | I25 | 241,895,316 | 125,060,174 | 23,871,213 | 7,864,240 | 16,006,973 | 32.94 | 19.09 | 6.62 | | | |
| Jomon | I27 | 241,450,064 | 126,487,721 | 22,510,933 | 6,543,998 | 15,966,935 | 29.07 | 17.80 | 6.61 | | | |
| Jomon | I9 | 10,988,506 | 5,355,997 | 163,147 | 607 | 162,540 | 0.37 | 3.05 | 1.48 | Petrous bone | NEBNext Ultra DNA library kit | Kyushu University, 24 Aug., 2016 |
| Jomon | I10 | 11,869,178 | 6,063,566 | 992,308 | 8,761 | 983,547 | 0.88 | 16.37 | 8.29 | | | |
| Jomon | Jomon_1 | 52,887,549 | 50,856,175 | 582,006 | 6,196 | 575,810 | 1.06 | 1.14 | 1.09 | | | |
| Jomon | Jomon_2 | 36,617,606 | 34,413,362 | 1,173,038 | 15,946 | 1,157,092 | 1.36 | 3.41 | 3.16 | | | |
| Jomon | Jomon_3 | 48,563,648 | 47,834,030 | 5,590,451 | 113,727 | 5,476,724 | 2.03 | 11.69 | 11.28 | Petrous bone | Allentoft et al. 2015 | CGG, Copenhagen, Oct., 2016 |
| Jomon | Jomon_4 | 58,511,078 | 56,245,786 | 4,845,321 | 88,276 | 4,757,045 | 1.82 | 8.61 | 8.13 | | | |
| Jomon | Jomon_5 | 43,866,323 | 42,562,678 | 2,946,263 | 386,321 | 2,559,942 | 13.11 | 6.92 | 5.84 | | | |

### Table S3

| Target | Ref 1 | Ref 2 | Amplitude | | | Decay | | |
|---|---|---|---|---|---|---|---|---|
| | | | Estimate | SE | Z | Estimate | SE | Z |
| Japanese | IK002 | Han | 0.00037 | 0.00006 | 6.3 | 49.8 | 7.6 | 6.6 |
| Ulchi | IK002 | Han | 0.00014 | 0.00005 | 2.7 | 20.8 | 12.6 | 1.7 |
| Ainu | IK002 | Japanese | 0.00018 | 0.00003 | 5.8 | 3.3 | 1.1 | 3.0 |

### Table S4

| W | X | Y | Z | D_score | D_sd | D_2sd | Z_score | SNPs |
|---|---|---|---|---|---|---|---|---|
| Mbuti | MA1 | Atayal | Ami | 0.0033 | 0.0050 | 0.0100 | 0.662 | 1285783 |
| Mbuti | MA1 | Burmese | Ami | -0.0012 | 0.0040 | 0.0081 | -0.287 | 1352503 |
| Mbuti | MA1 | Cambodian | Ami | 0.0021 | 0.0040 | 0.0081 | 0.528 | 1355903 |
| Mbuti | MA1 | Dai | Ami | -0.0008 | 0.0037 | 0.0074 | -0.209 | 1360951 |
| Mbuti | MA1 | Daur | Ami | -0.0070 | 0.0048 | 0.0095 | -1.467 | 1290590 |
| Mbuti | MA1 | Dusun | Ami | 0.0081 | 0.0041 | 0.0083 | 1.962 | 1354965 |
| Mbuti | MA1 | Han | Ami | -0.0010 | 0.0039 | 0.0078 | -0.252 | 1355943 |
| Mbuti | MA1 | Hezhen | Ami | -0.0009 | 0.0041 | 0.0083 | -0.210 | 1356366 |
| Mbuti | MA1 | Igorot | Ami | 0.0033 | 0.0038 | 0.0077 | 0.862 | 1354278 |
| Mbuti | MA1 | Japanese | Ami | 0.0008 | 0.0037 | 0.0073 | 0.225 | 1360859 |
| Mbuti | MA1 | Kinh | Ami | -0.0001 | 0.0040 | 0.0080 | -0.013 | 1355428 |
| Mbuti | MA1 | Korean | Ami | -0.0009 | 0.0041 | 0.0082 | -0.211 | 1353161 |
| Mbuti | MA1 | Lahu | Ami | -0.0007 | 0.0041 | 0.0082 | -0.174 | 1356419 |
| Mbuti | MA1 | Miao | Ami | 0.0024 | 0.0040 | 0.0080 | 0.590 | 1352492 |
| Mbuti | MA1 | Mongola | Ami | -0.0042 | 0.0040 | 0.0079 | -1.059 | 1353167 |
| Mbuti | MA1 | Naxi | Ami | -0.0013 | 0.0038 | 0.0075 | -0.346 | 1360569 |
| Mbuti | MA1 | Oroqen | Ami | -0.0136 | 0.0041 | 0.0082 | -3.311 | 1355451 |
| Mbuti | MA1 | She | Ami | -0.0009 | 0.0041 | 0.0083 | -0.211 | 1349530 |
| Mbuti | MA1 | Thai | Ami | 0.0017 | 0.0039 | 0.0078 | 0.439 | 1353479 |
| Mbuti | MA1 | Tu | Ami | -0.0039 | 0.0041 | 0.0083 | -0.952 | 1354164 |
| Mbuti | MA1 | Tujia | Ami | 0.0001 | 0.0040 | 0.0080 | 0.034 | 1350557 |
| Mbuti | MA1 | Xibo | Ami | -0.0003 | 0.0039 | 0.0079 | -0.065 | 1351587 |
| Mbuti | MA1 | Yi | Ami | 0.0001 | 0.0039 | 0.0079 | 0.029 | 1344798 |
| Mbuti | MA1 | Buryat | Ami | -0.0197 | 0.0043 | 0.0086 | -4.564 | 1339868 |
| Mbuti | MA1 | Chaplin | Ami | -0.0466 | 0.0052 | 0.0104 | -8.922 | 1302308 |
| Mbuti | MA1 | Naukan | Ami | -0.0500 | 0.0044 | 0.0088 | -11.332 | 1351888 |
| Mbuti | MA1 | Sireniki | Ami | -0.0484 | 0.0044 | 0.0088 | -10.961 | 1362400 |
| Mbuti | MA1 | Yupik | Ami | -0.0552 | 0.0048 | 0.0097 | -11.404 | 1020542 |
| Mbuti | MA1 | Even | Ami | -0.0194 | 0.0036 | 0.0073 | -5.323 | 1360314 |
| Mbuti | MA1 | Evenk | Ami | -0.0260 | 0.0048 | 0.0096 | -5.428 | 1305016 |
| Mbuti | MA1 | Itelman | Ami | -0.0498 | 0.0050 | 0.0099 | -10.047 | 1283108 |
| Mbuti | MA1 | Ket | Ami | -0.0550 | 0.0045 | 0.0091 | -12.116 | 1304859 |
| Mbuti | MA1 | Khanty | Ami | -0.0545 | 0.0040 | 0.0080 | -13.656 | 1361167 |
| Mbuti | MA1 | KomiIzhma | Ami | -0.0474 | 0.0046 | 0.0093 | -10.241 | 1358004 |
| Mbuti | MA1 | KomiObyachevo | Ami | -0.0520 | 0.0047 | 0.0094 | -11.120 | 1353998 |
| Mbuti | MA1 | Koryak | Ami | -0.0453 | 0.0045 | 0.0090 | -10.010 | 1339133 |
| Mbuti | MA1 | Ulchi | Ami | -0.0132 | 0.0042 | 0.0084 | -3.144 | 1354218 |
| Mbuti | MA1 | Nivkh | Ami | -0.0163 | 0.0043 | 0.0087 | -3.754 | 1343844 |
| Mbuti | MA1 | Chukchi | Ami | -0.0437 | 0.0049 | 0.0098 | -8.948 | 1300660 |
| Mbuti | MA1 | Chokhopani | Ami | -0.0037 | 0.0051 | 0.0102 | -0.724 | 1353283 |
| Mbuti | MA1 | Devilscave | Ami | 0.0091 | 0.0041 | 0.0082 | 2.213 | 1362226 |
| Mbuti | MA1 | IK002 | Ami | -0.0009 | 0.0054 | 0.0109 | -0.157 | 1124703 |

### Table S5

| W | X | Y | Z | D_score | D_sd | D_2sd | Z_score | SNPs |
|---|---|---|---|---|---|---|---|---|
| Mbuti | Atayal | MA1 | Ami | 0.1851 | 0.0053 | 0.0106 | 35.069 | 1285783 |
| Mbuti | Burmese | MA1 | Ami | 0.1230 | 0.0045 | 0.0090 | 27.428 | 1352503 |
| Mbuti | Cambodian | MA1 | Ami | 0.1304 | 0.0044 | 0.0088 | 29.484 | 1355903 |
| Mbuti | Dai | MA1 | Ami | 0.1527 | 0.0043 | 0.0086 | 35.418 | 1360951 |
| Mbuti | Daur | MA1 | Ami | 0.1247 | 0.0054 | 0.0107 | 23.267 | 1290590 |
| Mbuti | Dusun | MA1 | Ami | 0.1732 | 0.0046 | 0.0091 | 37.992 | 1354965 |
| Mbuti | Han | MA1 | Ami | 0.1528 | 0.0047 | 0.0094 | 32.468 | 1355943 |
| Mbuti | Hezhen | MA1 | Ami | 0.1286 | 0.0045 | 0.0090 | 28.494 | 1356366 |
| Mbuti | Igorot | MA1 | Ami | 0.1859 | 0.0047 | 0.0094 | 39.746 | 1354278 |
| Mbuti | Japanese | MA1 | Ami | 0.1428 | 0.0043 | 0.0086 | 33.163 | 1360859 |
| Mbuti | Kinh | MA1 | Ami | 0.1530 | 0.0045 | 0.0090 | 33.950 | 1355428 |
| Mbuti | Korean | MA1 | Ami | 0.1436 | 0.0046 | 0.0091 | 31.505 | 1353161 |
| Mbuti | Lahu | MA1 | Ami | 0.1440 | 0.0045 | 0.0090 | 31.830 | 1356419 |
| Mbuti | Miao | MA1 | Ami | 0.1533 | 0.0045 | 0.0090 | 34.003 | 1352492 |
| Mbuti | Mongola | MA1 | Ami | 0.1289 | 0.0047 | 0.0093 | 27.675 | 1353167 |
| Mbuti | Naxi | MA1 | Ami | 0.1361 | 0.0045 | 0.0089 | 30.539 | 1360569 |
| Mbuti | Oroqen | MA1 | Ami | 0.1147 | 0.0047 | 0.0095 | 24.187 | 1355451 |
| Mbuti | She | MA1 | Ami | 0.1576 | 0.0047 | 0.0093 | 33.826 | 1349530 |
| Mbuti | Thai | MA1 | Ami | 0.1391 | 0.0045 | 0.0090 | 31.067 | 1353479 |
| Mbuti | Tu | MA1 | Ami | 0.1229 | 0.0046 | 0.0091 | 26.929 | 1354164 |
| Mbuti | Tujia | MA1 | Ami | 0.1537 | 0.0046 | 0.0092 | 33.380 | 1350557 |
| Mbuti | Xibo | MA1 | Ami | 0.1303 | 0.0044 | 0.0087 | 29.833 | 1351587 |
| Mbuti | Yi | MA1 | Ami | 0.1401 | 0.0044 | 0.0089 | 31.499 | 1344798 |
| Mbuti | Buryat | MA1 | Ami | 0.0969 | 0.0047 | 0.0094 | 20.631 | 1339868 |
| Mbuti | Eskimo_Chaplin | MA1 | Ami | 0.0635 | 0.0056 | 0.0113 | 11.291 | 1302308 |
| Mbuti | Eskimo_Naukan | MA1 | Ami | 0.0532 | 0.0048 | 0.0095 | 11.176 | 1351888 |
| Mbuti | Eskimo_Sireniki | MA1 | Ami | 0.0569 | 0.0045 | 0.0090 | 12.602 | 1362400 |
| Mbuti | Eskimo_Yupik | MA1 | Ami | 0.0568 | 0.0052 | 0.0104 | 10.890 | 1020542 |
| Mbuti | Even | MA1 | Ami | 0.0926 | 0.0044 | 0.0088 | 20.995 | 1360314 |
| Mbuti | Evenk | MA1 | Ami | 0.1046 | 0.0054 | 0.0109 | 19.224 | 1305016 |
| Mbuti | Itelman | MA1 | Ami | 0.0652 | 0.0057 | 0.0114 | 11.467 | 1283108 |
| Mbuti | Ket | MA1 | Ami | 0.0259 | 0.0048 | 0.0097 | 5.358 | 1304859 |
| Mbuti | Khanty | MA1 | Ami | 0.0108 | 0.0043 | 0.0086 | 2.511 | 1361167 |
| Mbuti | KomiIzhma | MA1 | Ami | -0.0334 | 0.0046 | 0.0091 | -7.311 | 1358004 |
| Mbuti | KomiObyachevo | MA1 | Ami | -0.0336 | 0.0047 | 0.0094 | -7.128 | 1353998 |
| Mbuti | Koryak | MA1 | Ami | 0.0780 | 0.0051 | 0.0101 | 15.395 | 1339133 |
| Mbuti | Ulchi | MA1 | Ami | 0.1169 | 0.0045 | 0.0090 | 25.907 | 1354218 |
| Mbuti | Nivkh | MA1 | Ami | 0.1230 | 0.0050 | 0.0100 | 24.560 | 1343844 |
| Mbuti | Chukchi | MA1 | Ami | 0.0099 | 0.0053 | 0.0106 | 1.861 | 1300660 |
| Mbuti | Chokhopani | MA1 | Ami | 0.1137 | 0.0054 | 0.0108 | 21.098 | 1353283 |
| Mbuti | Devilscave | MA1 | Ami | 0.1369 | 0.0046 | 0.0092 | 29.867 | 1362226 |
| Mbuti | IK002 | MA1 | Ami | 0.1021 | 0.0059 | 0.0117 | 17.388 | 1124703 |

### Table S6

| W | X | Y | Z | D_score | D_sd | D_2sig | Z_score | SNPs |
|---|---|---|---|---|---|---|---|---|
| Mbuti | Ami | Atayal | MA1 | -0.1819 | 0.0052 | 0.0104 | -34.962 | 1285783 |
| Mbuti | Ami | Burmese | MA1 | -0.1241 | 0.0045 | 0.0089 | -27.738 | 1352503 |
| Mbuti | Ami | Cambodian | MA1 | -0.1283 | 0.0048 | 0.0095 | -26.966 | 1355903 |
| Mbuti | Ami | Dai | MA1 | -0.1534 | 0.0044 | 0.0088 | -34.851 | 1360951 |
| Mbuti | Ami | Daur | MA1 | -0.1316 | 0.0052 | 0.0104 | -25.420 | 1290590 |
| Mbuti | Ami | Dusun | MA1 | -0.1654 | 0.0045 | 0.0091 | -36.444 | 1354965 |
| Mbuti | Ami | Han | MA1 | -0.1537 | 0.0046 | 0.0093 | -33.161 | 1355943 |
| Mbuti | Ami | Hezhen | MA1 | -0.1295 | 0.0048 | 0.0095 | -27.205 | 1356366 |
| Mbuti | Ami | Igorot | MA1 | -0.1827 | 0.0045 | 0.0089 | -40.967 | 1354278 |
| Mbuti | Ami | Japanese | MA1 | -0.1420 | 0.0045 | 0.0090 | -31.558 | 1360859 |
| Mbuti | Ami | Kinh | MA1 | -0.1531 | 0.0046 | 0.0092 | -33.256 | 1355428 |
| Mbuti | Ami | Korean | MA1 | -0.1445 | 0.0047 | 0.0093 | -30.972 | 1353161 |
| Mbuti | Ami | Lahu | MA1 | -0.1447 | 0.0045 | 0.0090 | -32.191 | 1356419 |
| Mbuti | Ami | Miao | MA1 | -0.1510 | 0.0045 | 0.0090 | -33.408 | 1352492 |
| Mbuti | Ami | Mongola | MA1 | -0.1330 | 0.0046 | 0.0092 | -29.041 | 1353167 |
| Mbuti | Ami | Naxi | MA1 | -0.1373 | 0.0046 | 0.0091 | -30.171 | 1360569 |
| Mbuti | Ami | Oroqen | MA1 | -0.1282 | 0.0047 | 0.0093 | -27.479 | 1355451 |
| Mbuti | Ami | She | MA1 | -0.1585 | 0.0046 | 0.0093 | -34.224 | 1349530 |
| Mbuti | Ami | Thai | MA1 | -0.1374 | 0.0044 | 0.0088 | -31.082 | 1353479 |
| Mbuti | Ami | Tu | MA1 | -0.1268 | 0.0048 | 0.0095 | -26.567 | 1354164 |
| Mbuti | Ami | Tujia | MA1 | -0.1535 | 0.0046 | 0.0092 | -33.208 | 1350557 |
| Mbuti | Ami | Xibo | MA1 | -0.1306 | 0.0046 | 0.0092 | -28.527 | 1351587 |
| Mbuti | Ami | Yi | MA1 | -0.1400 | 0.0046 | 0.0091 | -30.753 | 1344798 |
| Mbuti | Ami | Buryat | MA1 | -0.1164 | 0.0048 | 0.0096 | -24.375 | 1339868 |
| Mbuti | Ami | Chaplin | MA1 | -0.1098 | 0.0053 | 0.0106 | -20.766 | 1302308 |
| Mbuti | Ami | Naukan | MA1 | -0.1029 | 0.0048 | 0.0096 | -21.541 | 1351888 |
| Mbuti | Ami | Sireniki | MA1 | -0.1051 | 0.0047 | 0.0094 | -22.393 | 1362400 |
| Mbuti | Ami | Yupik | MA1 | -0.1117 | 0.0051 | 0.0103 | -21.723 | 1020542 |
| Mbuti | Ami | Even | MA1 | -0.1118 | 0.0045 | 0.0090 | -24.888 | 1360314 |
| Mbuti | Ami | Evenk | MA1 | -0.1302 | 0.0054 | 0.0108 | -24.097 | 1305016 |
| Mbuti | Ami | Itelman | MA1 | -0.1146 | 0.0052 | 0.0104 | -22.125 | 1283108 |
| Mbuti | Ami | Ket | MA1 | -0.0808 | 0.0048 | 0.0096 | -16.885 | 1304859 |
| Mbuti | Ami | Khanty | MA1 | -0.0652 | 0.0043 | 0.0086 | -15.090 | 1361167 |
| Mbuti | Ami | KomiIzhma | MA1 | -0.0140 | 0.0047 | 0.0093 | -3.002 | 1358004 |
| Mbuti | Ami | KomiObyachevo | MA1 | -0.0185 | 0.0046 | 0.0092 | -3.995 | 1353998 |
| Mbuti | Ami | Koryak | MA1 | -0.1228 | 0.0048 | 0.0097 | -25.435 | 1339133 |
| Mbuti | Ami | Ulchi | MA1 | -0.1299 | 0.0047 | 0.0093 | -27.806 | 1354218 |
| Mbuti | Ami | Nivkh | MA1 | -0.1390 | 0.0048 | 0.0095 | -29.134 | 1343844 |
| Mbuti | Ami | Chukuchi | MA1 | -0.0536 | 0.0051 | 0.0102 | -10.561 | 1300660 |
| Mbuti | Ami | Chokhopani | MA1 | -0.1174 | 0.0051 | 0.0103 | -22.821 | 1353283 |
| Mbuti | Ami | Devilscave | MA1 | -0.1279 | 0.0047 | 0.0094 | -27.119 | 1362226 |
| Mbuti | Ami | IK002 | MA1 | -0.1030 | 0.0053 | 0.0106 | -19.350 | 1124703 |

Fig. S1

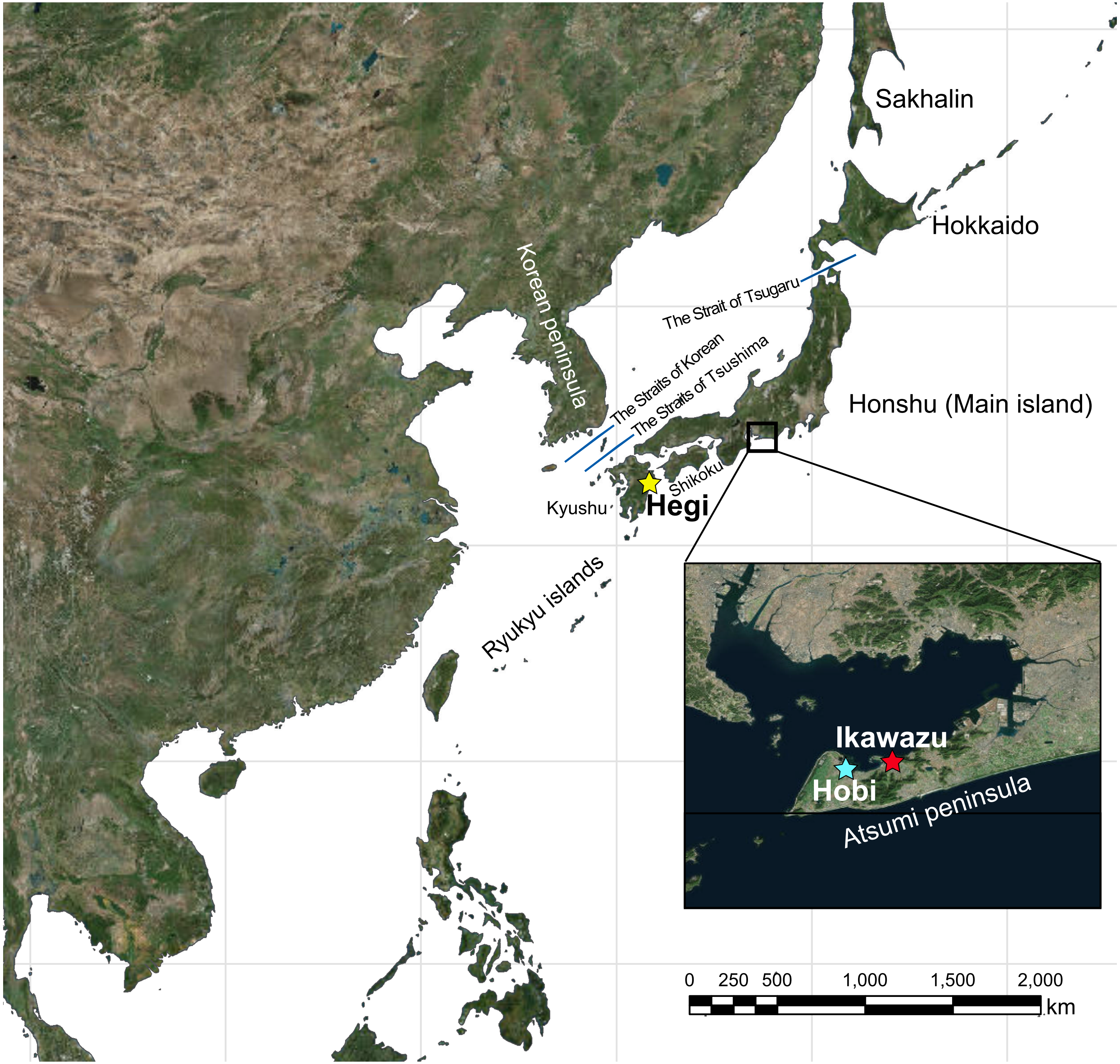

Sakhalin
Hokkaido
Korean peninsula
The Strait of Tsugaru
The Straits of Korean
The Straits of Tsushima
Honshu (Main island)
Shikoku
Kyushu
Hegi
Ryukyu islands
Ikawazu
Hobi
Atsumi peninsula
0 250 500 1,000 1,500 2,000
km

Fig. S2

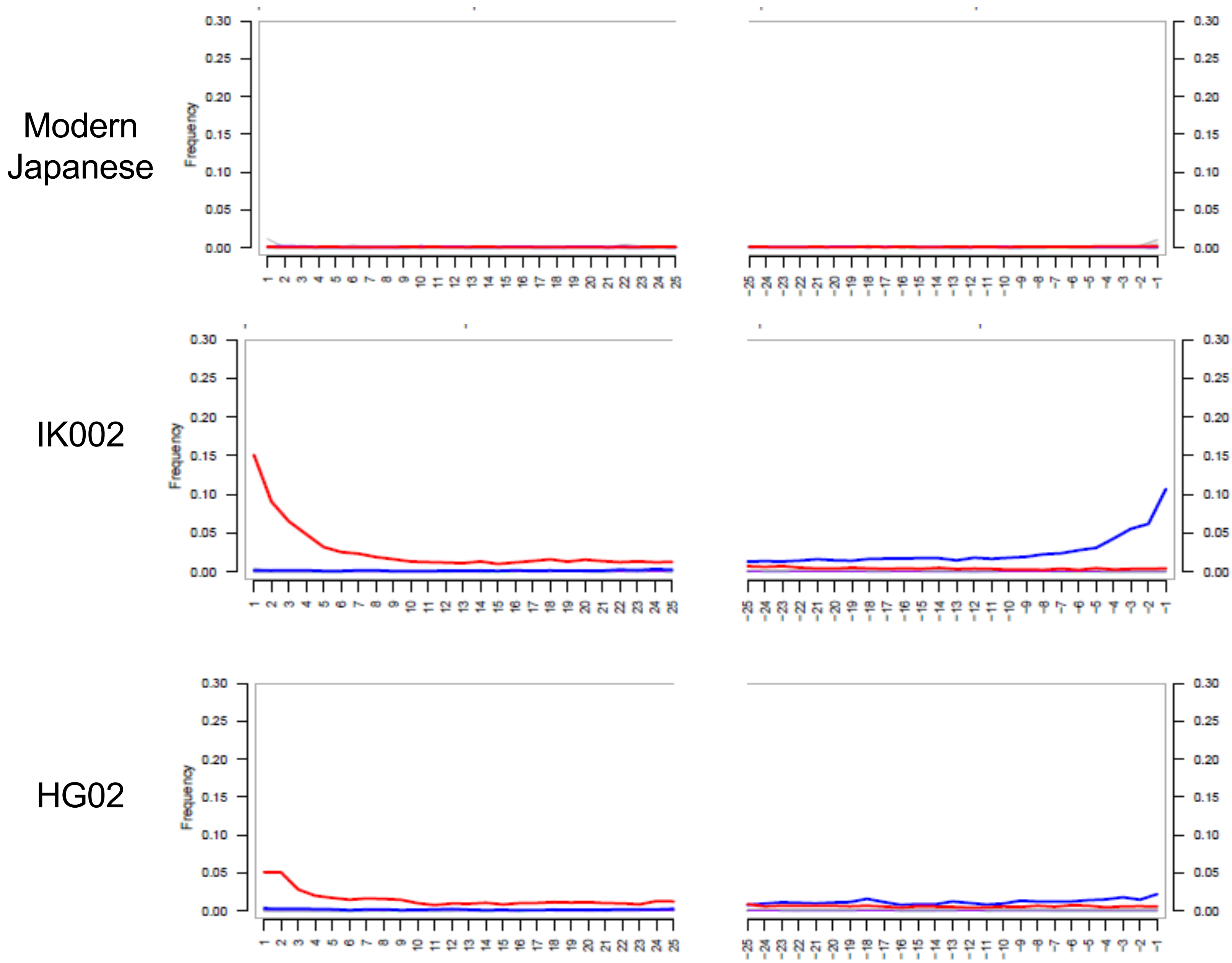

Modern Japanese
IK002
HG02
Frequency
0.00
0.05
0.10
0.15
0.20
0.25
0.30

Fig. S3

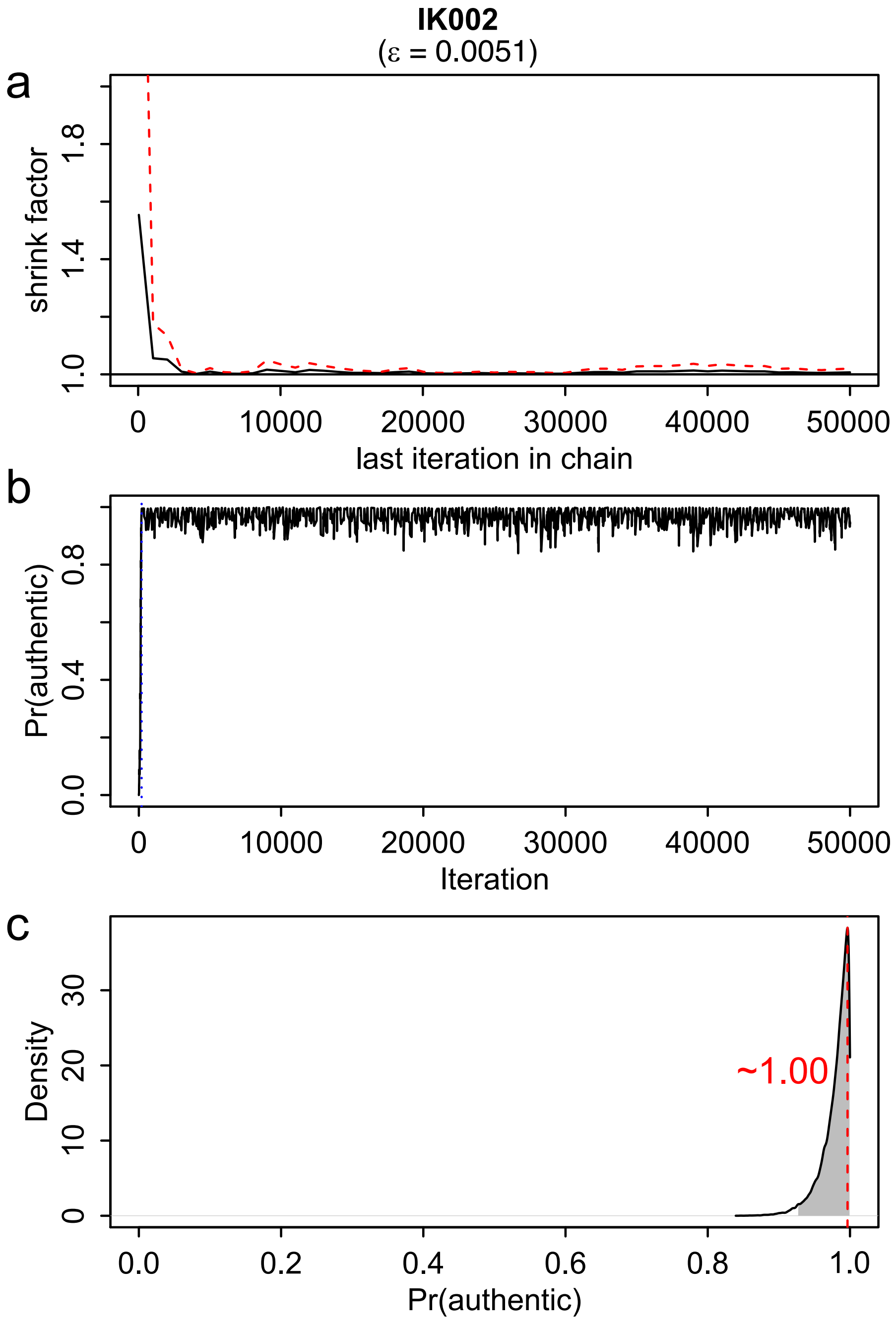

IK002
(ε = 0.0051)
a
shrink factor
1.0
1.4
1.8
0
10000
20000
30000
40000
50000
last iteration in chain
b
Pr(authentic)
0.0
0.4
0.8
0
10000
20000
30000
40000
50000
Iteration
c
Density
0
10
20
30
~1.00
0.0
0.2
0.4
0.6
0.8
1.0
Pr(authentic)

Fig. S4

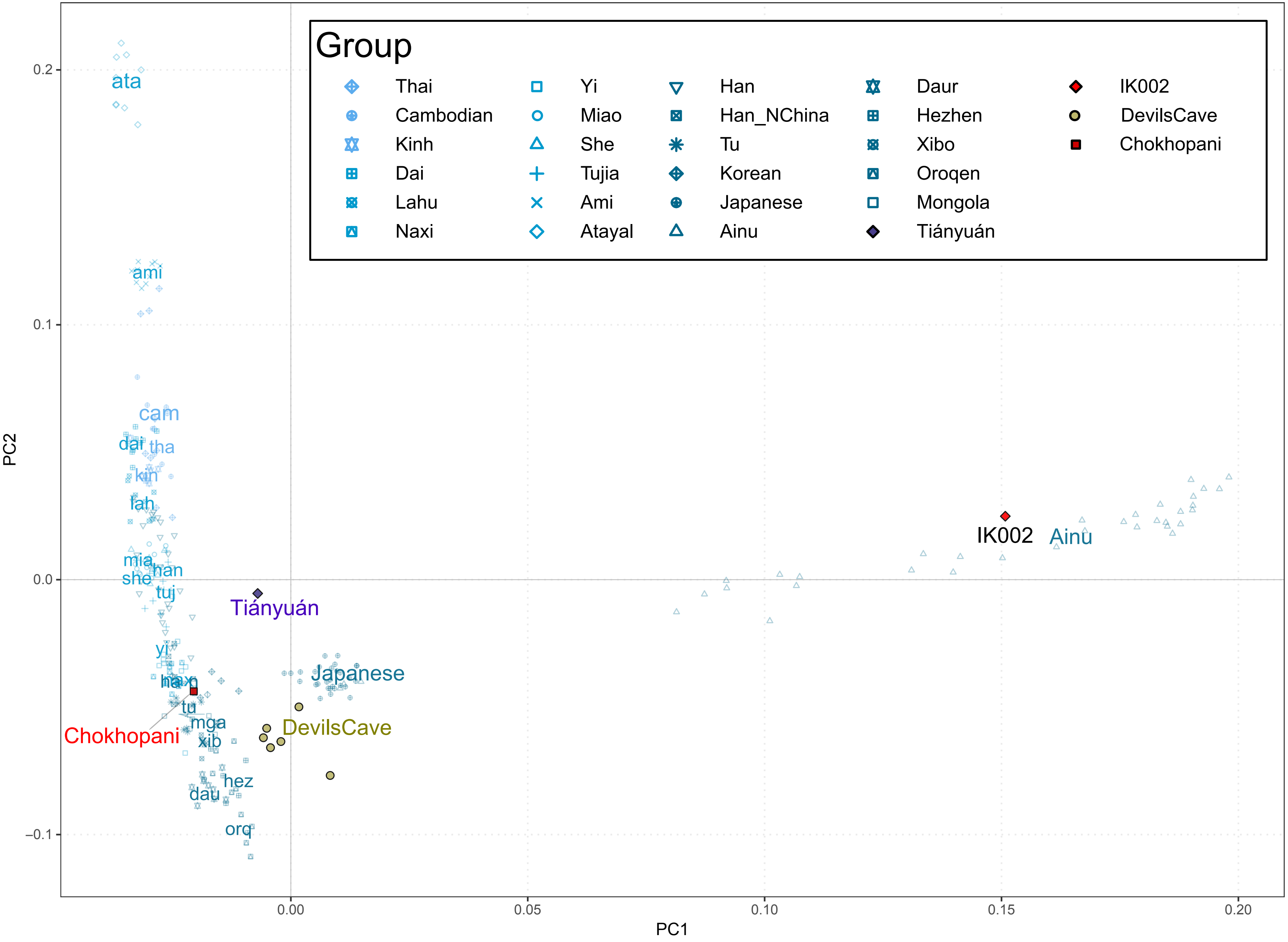
Group
Thai
Cambodian
Kinh
Dai
Lahu
Naxi
Yi
Miao
She
Tujia
Ami
Atayal
Han
Han_NChina
Tu
Korean
Japanese
Ainu
Daur
Hezhen
Xibo
Oroqen
Mongola
Tiányuán
IK002
DevilsCave
Chokhopani
ata
ami
cam
dai
tha
kin
lah
mia
han
she
tuj
Tiányuán
yi
Japanese
tu
mga
xib
Chokhopani
DevilsCave
hez
dau
orq
IK002
Ainu
PC2
PC1
0.2
0.1
0.0
−0.1
0.00
0.05
0.10
0.15
0.20

Fig. S5

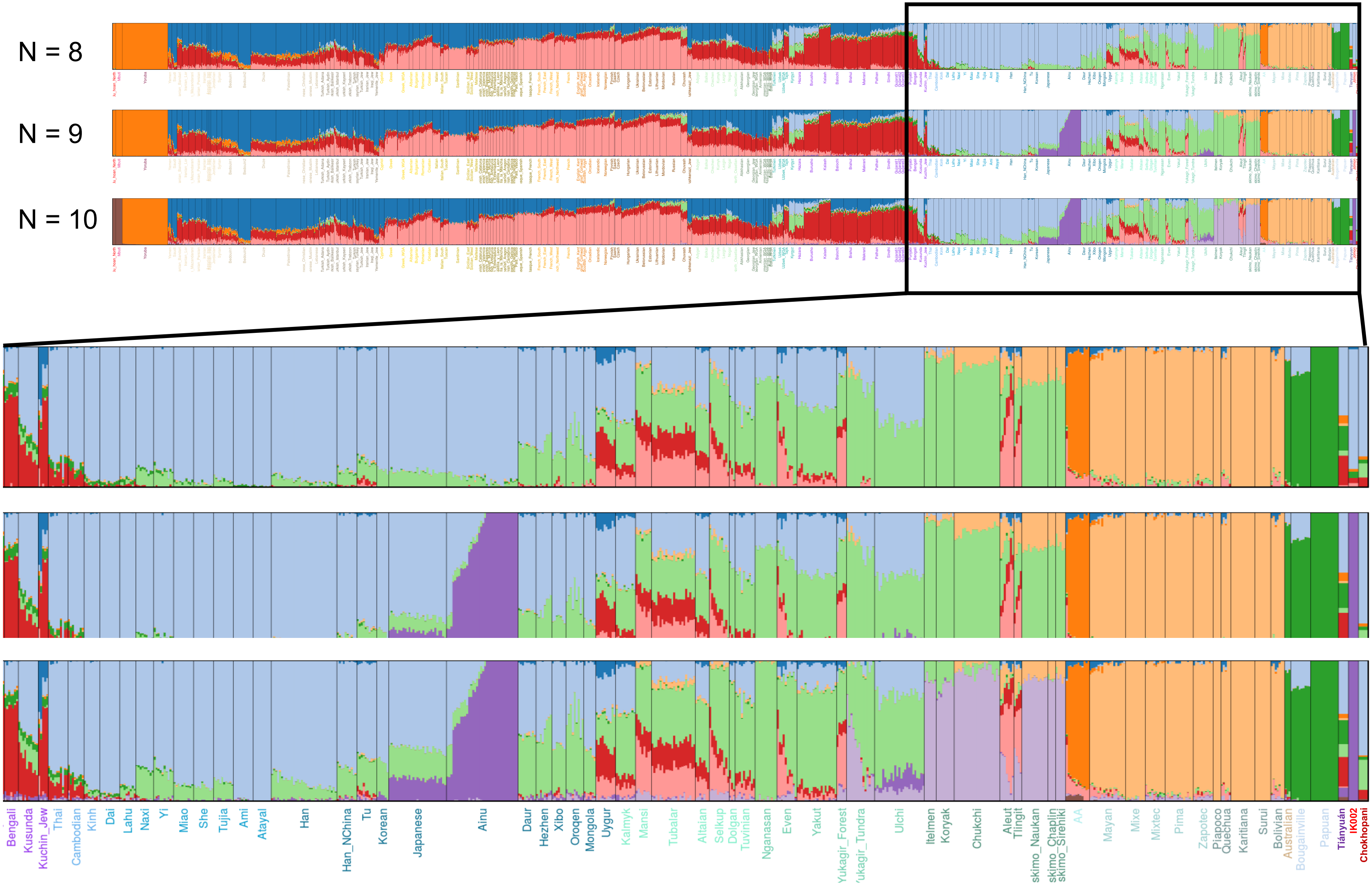
N = 8
N = 9
N = 10
Bengali
Kusunda
Kuchin_Jew
Thai
Cambodian
Kinh
Dai
Lahu
Naxi
Yi
Miao
She
Tujia
Ami
Atayal
Han
Han_NChina
Tu
Korean
Japanese
Ainu
Daur
Hezhen
Xibo
Oroqen
Mongola
Uygur
Kalmyk
Mansi
Tubalar
Altaian
Selkup
Dolgan
Tuvinian
Nganasan
Even
Yakut
Yukagir_Forest
'ukagir_Tundra
Ulchi
Itelmen
Koryak
Chukchi
Aleut
Tlingit
skimo_Naukan
skimo_Chaplin
skimo_Sireniki
AA
Mayan
Mixe
Mixtec
Pima
Zapotec
Piapoco
Quechua
Karitiana
Surui
Bolivian
Australian
Bougainville
Papuan
Tiányuán
IK002
Chokhopani

Fig. S6

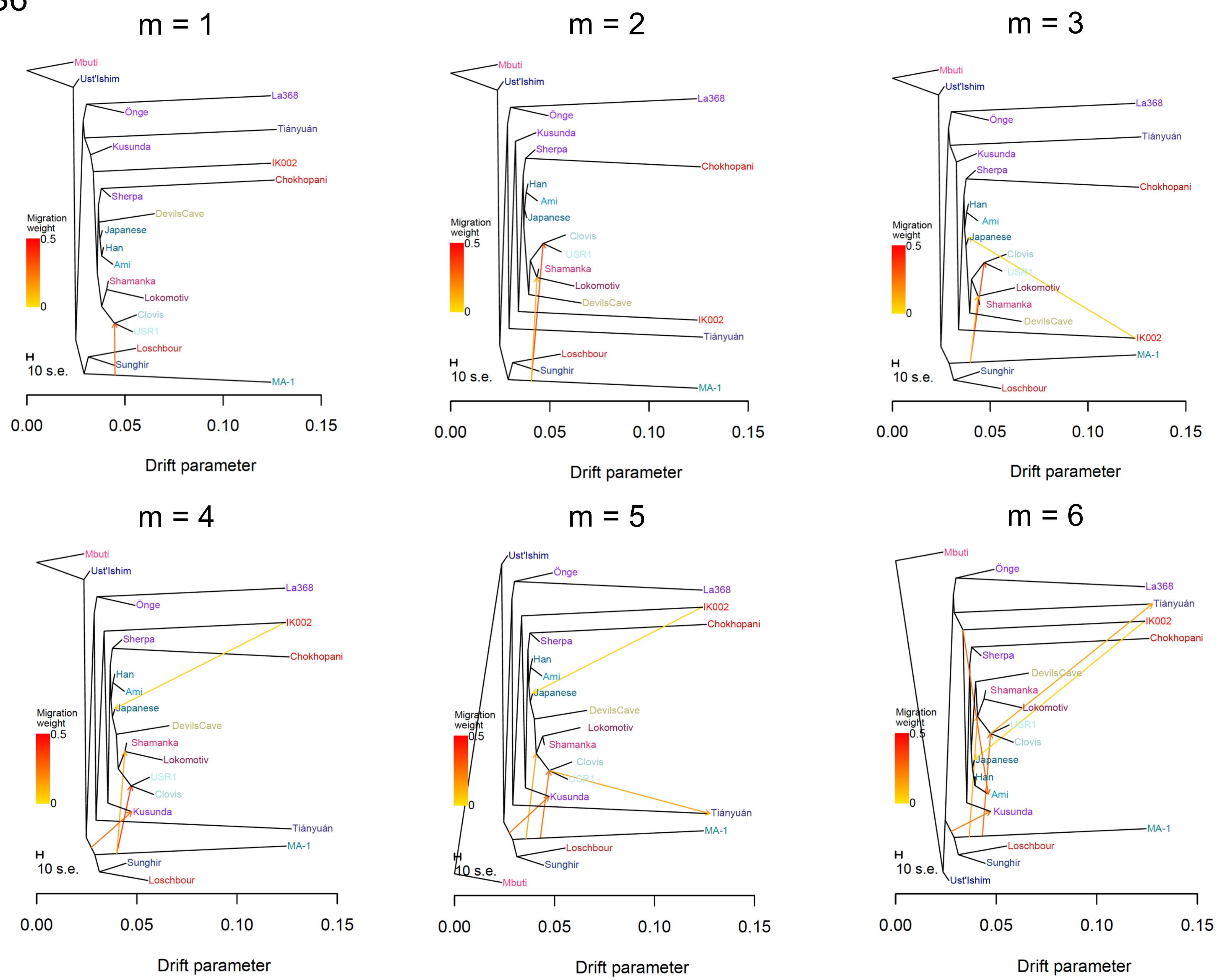

m = 1
Mbuti
Ust'Ishim
La368
Önge
Tiányuán
Kusunda
IK002
Chokhopani
Sherpa
DevilsCave
Japanese
Han
Ami
Shamanka
Lokomotiv
Clovis
USR1
Loschbour
Sunghir
MA-1
Migration weight
0.5
0
10 s.e.
0.00
0.05
0.10
0.15
Drift parameter
m = 2
m = 3
m = 4
m = 5
m = 6

Fig. S7

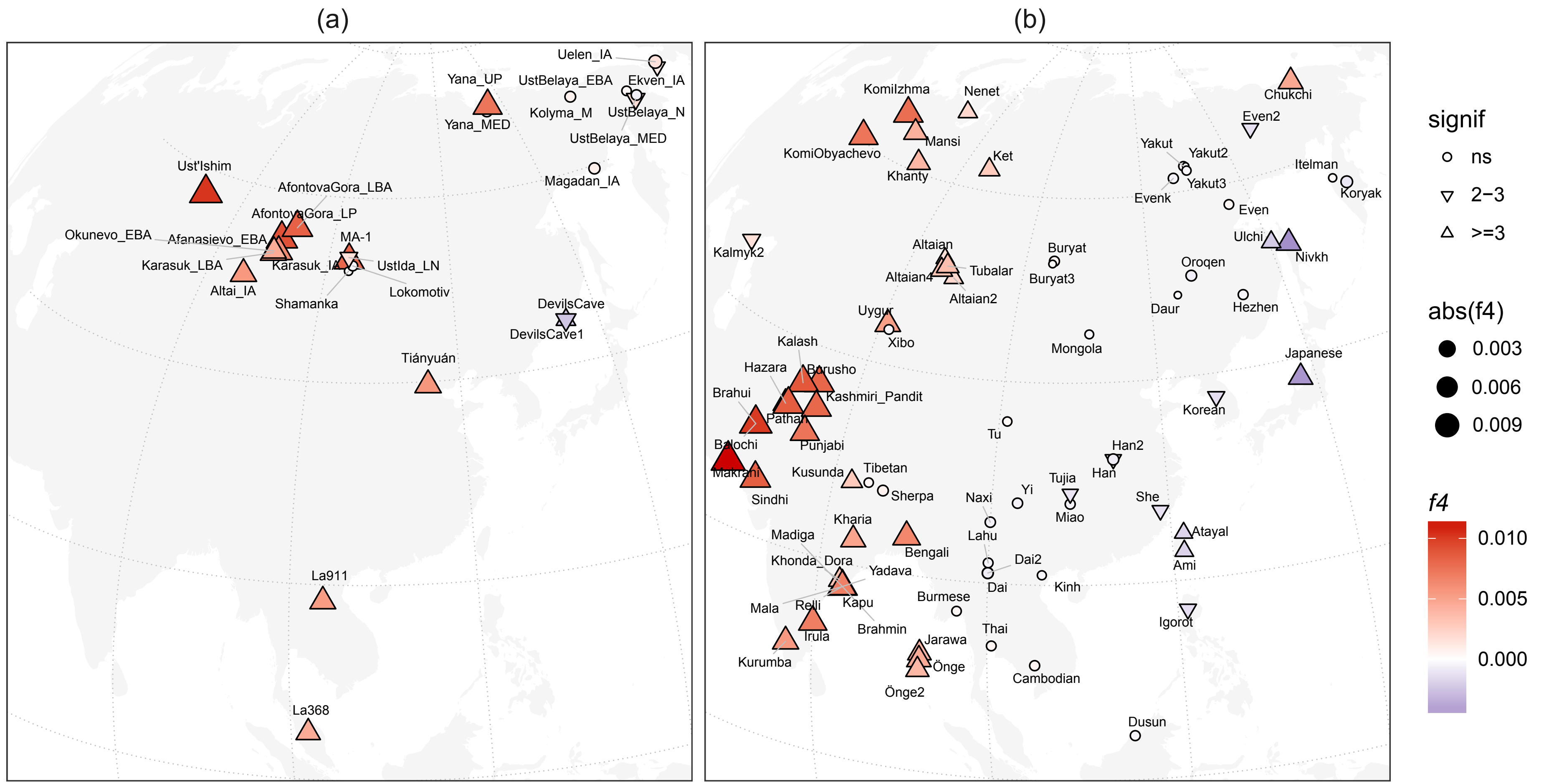
(a)
(b)
Uelen_IA
Yana_UP
UstBelaya_EBA
Ekven_IA
Kolyma_M
UstBelaya_N
Yana_MED
UstBelaya_MED
Magadan_IA
Ust'Ishim
AfontovaGora_LBA
AfontovaGora_LP
Okunevo_EBA
Afanasievo_EBA
MA-1
Karasuk_LBA
Karasuk_IA
UstIda_LN
Altai_IA
Lokomotiv
Shamanka
DevilsCave
DevilsCave1
Tiányuán
La911
La368
Komilzhma
Nenet
Chukchi
Even2
KomiObyachevo
Mansi
Ket
Yakut
Yakut2
Khanty
Itelman
Evenk
Yakut3
Koryak
Even
Kalmyk2
Altaian
Buryat
Ulchi
Nivkh
Tubalar
Oroqen
Altaian4
Buryat3
Altaian2
Daur
Hezhen
Uygur
Xibo
Mongola
Kalash
Japanese
Hazara
Burusho
Brahui
Kashmiri_Pandit
Korean
Pathan
Tu
Balochi
Punjabi
Han2
Makrani
Tibetan
Han
Kusunda
Tujia
Sindhi
Sherpa
Yi
Naxi
She
Miao
Kharia
Atayal
Madiga
Lahu
Bengali
Ami
Khonda_Dora
Dai2
Yadava
Dai
Kinh
Mala
Relli
Kapu
Burmese
Igorot
Irula
Brahmin
Thai
Jarawa
Kurumba
Önge
Cambodian
Önge2
Dusun
signif
ns
2−3
>=3
abs(f4)
0.003
0.006
0.009
f4
0.010
0.005
0.000

Fig. S8

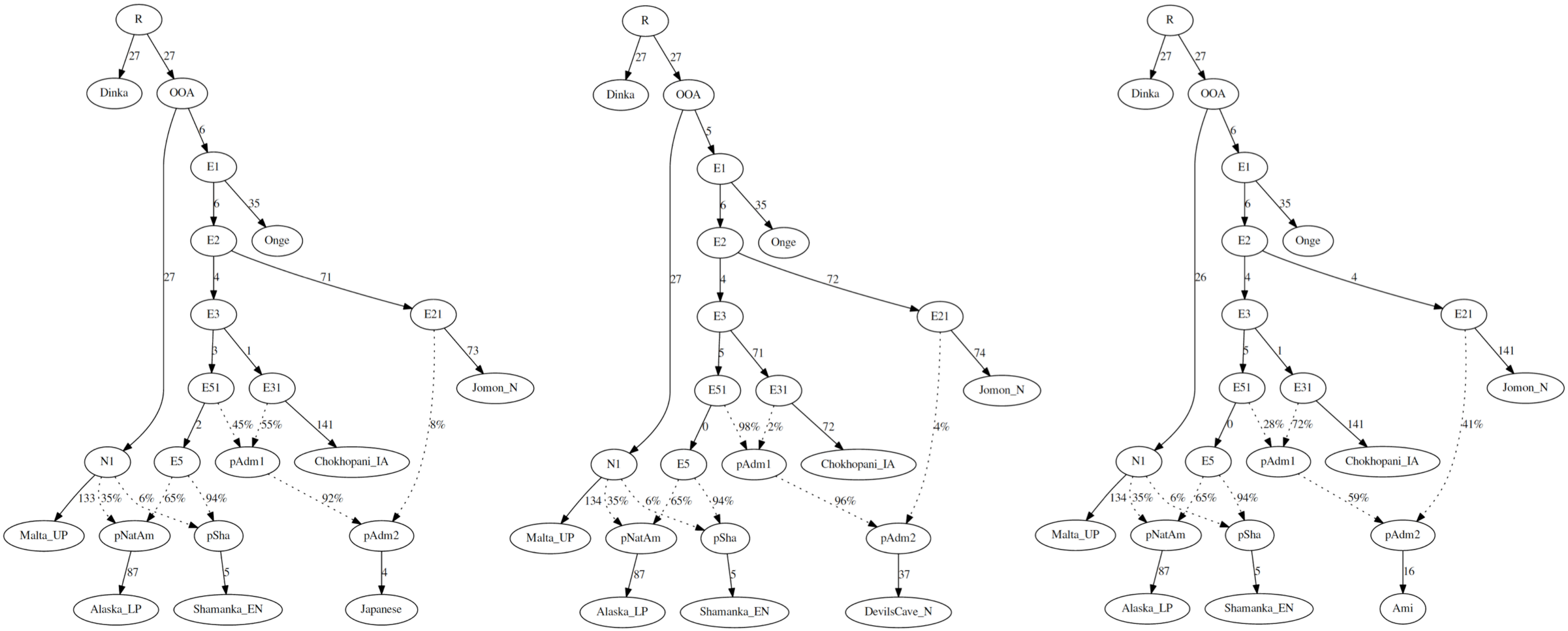

R
27
27
Dinka
OOA
6
27
E1
6
35
E2
Onge
4
71
E3
E21
3
1
73
E51
E31
Jomon_N
2
45%
55%
141
8%
N1
E5
pAdm1
Chokhopani_IA
133
35%
6%
65%
94%
92%
Malta_UP
pNatAm
pSha
pAdm2
87
5
4
Alaska_LP
Shamanka_EN
Japanese
R
27
27
Dinka
OOA
5
27
E1
6
35
E2
Onge
4
72
E3
E21
5
71
74
E51
E31
Jomon_N
0
98%
2%
72
4%
N1
E5
pAdm1
Chokhopani_IA
134
35%
6%
65%
94%
96%
Malta_UP
pNatAm
pSha
pAdm2
87
5
37
Alaska_LP
Shamanka_EN
DevilsCave_N
R
27
27
Dinka
OOA
6
26
E1
6
35
E2
Onge
4
4
E3
E21
5
1
141
E51
E31
Jomon_N
0
28%
72%
141
41%
N1
E5
pAdm1
Chokhopani_IA
134
35%
6%
65%
94%
59%
Malta_UP
pNatAm
pSha
pAdm2
87
5
16
Alaska_LP
Shamanka_EN
Ami
